# Label- and slide-free multispectral quantitative epi-illumination deep-UV microscopy

**DOI:** 10.64898/2026.01.12.699112

**Authors:** Mingxuan Si, Viswanath Gorti, Aaron D. Silva Trenkle, Arjun Renjith, Brienna E. Heinsz, Gabe A. Kwong, Francisco E. Robles

## Abstract

Label-free and slide-free imaging is highly desired in clinical pathology because it holds the potential to avoid time- and labor-intensive tissue processing and chemical staining while preserving molecular information for downstream analyses. Deep-ultraviolet (UV) microscopy offers high-resolution, label-free molecular contrast via short wavelengths and intrinsic biomolecular absorption, but prior implementations have been limited to the analysis of thin sections and cell monolayers. Here, we present a fast, low-cost, LED-based, epi-illumination deep-UV microscope (epi-DUV) for label- and slide-free imaging of fresh, thick tissues. Using 255 nm and 280 nm absorption images, and tryptophan autofluorescence, the method yields quantitative maps of nucleic acid mass, protein mass, and quantum yield. Moreover, H&E-like contrast can be generated using native 255-nm absorption images. The system achieves 0.5 µm lateral resolution with an effective slice thickness of ∼6 µm across a 707 µm × 707 µm field of view and uses ∼330-ms exposure. To the best of our knowledge, this is the first demonstration of quantitative deep-UV molecular imaging of fresh, unlabeled thick tissues. Epi-DUV has significant potential to streamline the histopathology workflow while adding objective molecular readouts, enabling point-of-care assessment of unprocessed specimens (e.g., rapid intraoperative evaluation).

## Introduction

Formalin-fixed and paraffin-embedded (FFPE) tissue processing remains the gold standard for histopathological examination, enabling diagnosis of a variety of diseases, including cancers, infections, and many other abnormalities (*1–3*). FFPE relies on fixing tissues, embedding in paraffin wax, preparing thin tissue sections mounted on glass slides, followed by staining (e.g., hematoxylin and eosin (H&E)) to reveal tissue morphology (and in some cases, molecular composition) with high contrast. While these techniques provide high diagnostic utility and have been a mainstay tool in clinical pathology, they are time- and labor-intensive, requiring specialized equipment, trained personnel, and several chemical processing steps. The workflow, from sample acquisition to pathological evaluation, can take many hours to days (*3–5*), limiting intraoperative or point-of-care decision-making which ultimately delays patient care. Although frozen sections provide a faster alternative to FFPE sections, with turnaround times of 20-30 mins, their diagnostic quality is significantly lower (*3*, *6*, *7*). Furthermore, these analyses cannot be performed in low-resource settings without access to histology laboratories and the required equipment and supplies.

Slide-free histology has recently emerged as a promising alternative, offering the potential for direct point-of-care imaging of intact tissues with no or minimal processing. This can enable real-time histological evaluation, reducing time to diagnosis while preserving tissue for downstream analyses. Several optical techniques have been applied for slide-free histology, including point scanning methods such as confocal microscopy (*8–11*) and nonlinear multiphoton microscopy (*12–15*), which require some tissue processing for fluorescence labelling. More advanced label-free methods have also been introduced, such as stimulated Raman scattering (*16–20*), and photoacoustic microscopy (*21–28*). These techniques provide 3D histological images of un- (or minimally-) processed tissues, but their clinical translation/adoption is hampered by the complexity, expense, and relatively large footprint of the systems. Wide-field imaging methods, including microscopy with ultraviolet surface excitation (MUSE) (*29*) and structured illumination microscopy (SIM) (*30*, *31*), have also been applied for slide-free imaging. These methods use much simpler instrumentation (i.e., lower costs and lower system complexity) with spatial multiplexing capabilities that offer faster acquisition times and a more compact system. However, these methods also require fluorescence labelling, which necessitates some degree of sample preparation. While fluorescent labeling is not as onerous a process as the entire FFPE and staining pipeline, it does introduce other caveats, including variable uptake of the fluorescent agent (particularly problematic for fresh tissues), and signal degradation via photobleaching. (Typically, tissues have to be fixed for optimal dye uptake, which adds to the processing time.) Furthermore, the very need for the fluorescent compounds and other necessary reagents can be a significant added expense when imaging large volumes of tissues. Therefore, it is clinically desirable to have an imaging modality that is not only slide-free, fast, compact and low cost but also label-free.

Deep-ultraviolet (UV) microscopy has recently emerged as a powerful approach for wide-field label-free molecular imaging (*32–40*). By leveraging the intrinsic absorption properties of biomolecules, such as nucleic acids and proteins, in the deep-UV region of the spectrum (200–300 nm), this technique enables high-resolution, label-free, wide-field molecular imaging with simple and low-cost instrumentation. Deep-UV microscopy has been recently demonstrated for hematology and histopathology applications, including label-free molecular analysis of prostate tissue sections (*34*, *35*). One of the key advantages of deep-UV microscopy is its ability to enable quantitative mapping of biomolecules, providing pixel-level estimates of nucleic acid and protein mass with femtogram sensitivity derived directly from the absorption properties of these biomolecules (*40*, *41*). By transforming qualitative images into quantitative maps, quantitative deep-UV microscopy has the potential to enable objective molecular phenotyping and real-time point-of-care assessment of the cellular and molecular composition, serving as an auxiliary readout for cytology and histology applications. However, prior implementations of deep-UV microscopy operate in transmission, hindering their applicability to unprocessed thick tissues.

Here, we present multispectral epi-illumination deep-UV microscopy (epi-DUV) to overcome the limitations of transmission-based deep-UV microscopy and existing slide-free methods. Epi-DUV enables rapid, label-free molecular imaging of fresh, unprocessed, thick tissues with subcellular resolution. By way of multiple scattering within the thick tissue in combination with the strong UV absorption, the acquired epi-DUV images effectively resemble transmission-based images of the tissue surface, which can be quantified to yield quantitative maps of nucleic acid mass and protein mass with femtogram sensitivity, as well as quantum yield. Moreover, images acquired with 255 nm illumination can readily be color-adjusted to yield H&E-like contrast. We demonstrate the utility of the approach by imaging fresh murine and human tissue samples. To the best of our knowledge, this is the first demonstration of quantitative deep-UV molecular imaging of fresh, thick tissues. Together, our system is fast, label-free, and applicable for a variety of histopathology applications, offering a powerful tool for quantitative, stain- and slide-free histology.

## Results

### Histological imaging with epi-illumination deep-UV (epi-DUV) imaging system

The epi-DUV system (Fig. 1A) is equipped with four narrow-band LEDs, two with a center wavelength of 255 nm and two at 280 nm. The LEDs are directed to illuminate the bottom of the sample (which is placed atop a quartz slide) through a fused-silica light rod that acts as a light guide (Fig. 1B, C). The LEDs from matching wavelengths and their respective light guides are oriented opposite to each other as shown in Fig. 1C (blue arrows for 255 nm LEDs’ light guides; purple arrows for 280 nm LEDs’ light guides). This orientation increases the power (6.3 mW at the sample) and illumination homogeneity at the sample.

**Fig. 1.**
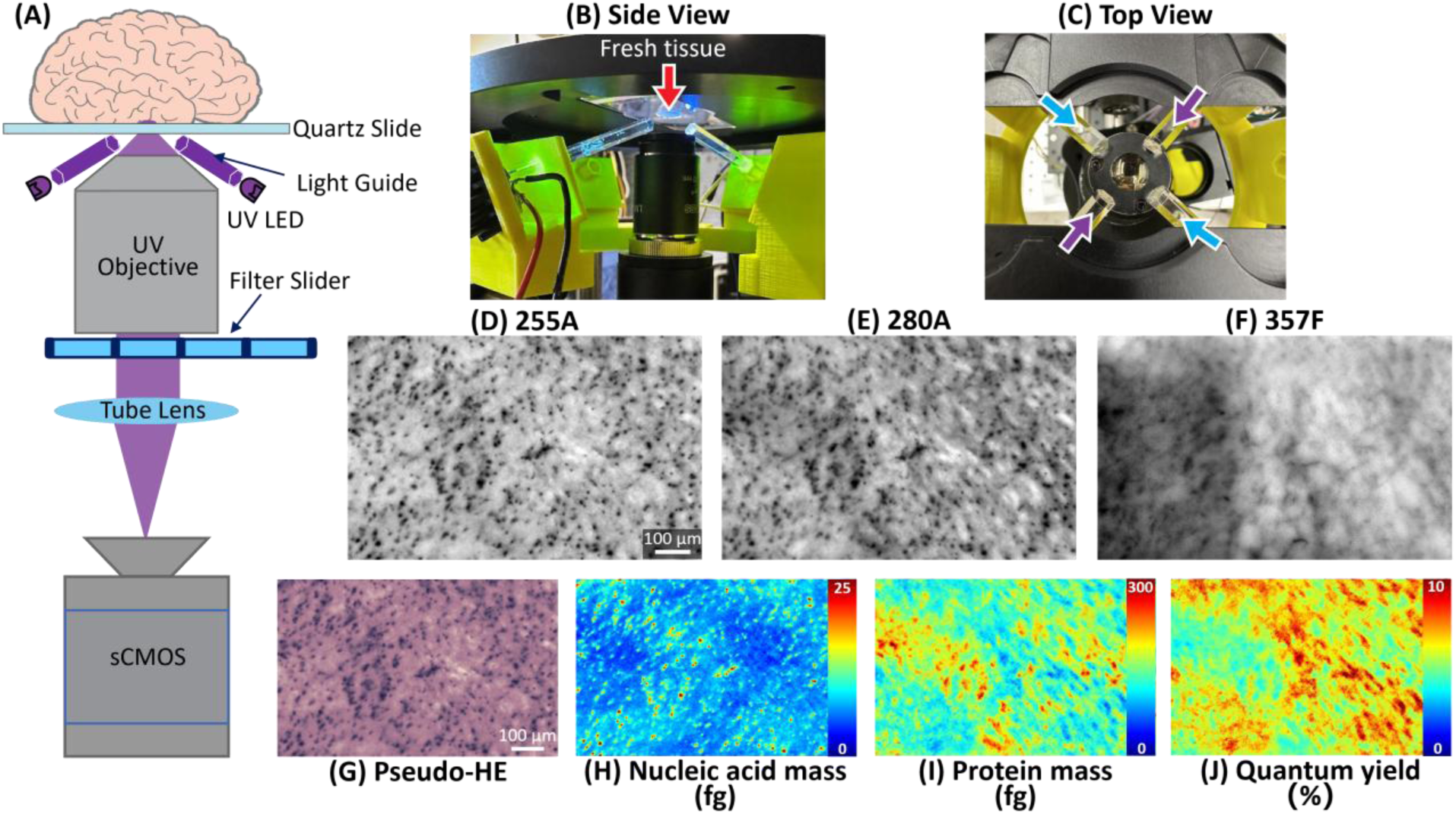
Epi-DUV imaging system setup and representative images from fresh mouse brain sample 1. (**A**) Schematic of the epi-DUV system, showing only two LEDs for better visualization. (**B**) Side view from just underneath the sample stage, showing only two LEDs for better visualization. Red arrow indicates the location of the fresh tissue. (**C**) Top view from above the sample stage. Blue arrows correspond to the 255 nm LEDs’ light guide. Purple arrows correspond to the 280 nm LEDs’ light guide. (**D**) Absorption image with 255 nm illumination and detection. (**E**) Absorption image with 280 nm illumination and detection. (**F**) Tryptophan autofluorescence image with 255 nm excitation and 357 nm detection. (**G**) Pseudo-H&E image of the same area using the 255A image only. (**H**) Nucleic acid mass map of the same area (in femtograms). (**I**) Protein mass map of the same area (in femtograms). (**J**) Tryptophan quantum yield map of the same area.

We generate image contrast based on either the tissue absorption or autofluorescence. We collect the 255 nm absorption (255A) and 280 nm absorption (280A) information by illuminating the sample with the two 255 nm LEDs and two 280 nm LEDs, respectively, and capturing the diffusely back-reflected light (*42*) from the bottom surface of the sample. To collect the two absorption images, we also use bandpass filters (10 nm spectral bandwidth) centered at 255 nm and 280 nm, respectively, to remove any autofluorescence. Note that 255 nm and 280 nm correspond to the absorption peaks of nucleic acids and proteins (*41*, *43*), thus the absorption images (Fig. 1D-E) provide direct insight into their spatial distribution. We also collect autofluorescence information using the same 255 nm illumination for excitation, which provides additional contrast for fluorescent molecules. Since UV autofluorescence is dominated by tryptophan (*44*, *45*), we incorporate a bandpass filter centered at 357 nm, which corresponds to the tryptophan emission peak— this configuration selectively capture tryptophan autofluorescence. Note that 255 nm is not at the excitation peak of tryptophan (*46*), but this wavelength is chosen for excitation to generate negative contrast of cell nuclei due to their strong absorption at this wavelength (Fig. 1F).

From the 255A image alone, we perform a fast, simple, and accurate “one-click” transformation to generate a realistic pseudo-H&E image (Fig. 1G) using simple Red, Green, and Blue (RGB) histogram matching between the 255A image and a sample H&E image of a similar region (see Materials and Methods). The generated pseudo-H&E images exhibit comparable nuclear contrast and density to real histological images, both of which are essential for pathologists in making accurate diagnoses.

To quantify molecular information (Fig. 1H–J), it is crucial to understand light-tissue interactions in the deep UV range. Tissue exhibits strong absorption due to the characteristic absorption peaks of nucleic acids and proteins (*40*, *41*). Additionally, the scattering behavior of tissue in this range is assumed to follow Lambertian (isotropic) scattering at a depth near the surface. This behavior arises from a small anisotropy factor (g∼< 0.5) and a large scattering coefficient in biological tissues (*47–50*). Under these conditions, it is reasonable to assume that a maximum of two scattering events occur before the scattering becomes fully isotropic.

We define the layer beneath the tissue surface where isotropic scattering occurs as the "diffusive layer". Within the space between this layer and the tissue surface, isotropically scattered photons propagate back through the tissue surface again while undergoing significant absorption. Photons that are not absorbed escape the tissue and are captured by the objective lens. We assume that the diffusive layer is so close to the tissue surface that light does not undergo an additional scattering event before exiting. In other words, we posit that the absorption coefficient in the deep UV range is comparable to, if not greater than, the scattering coefficient. This assumes that photons either escape the surface or are absorbed before contributing to the signal. These scattered photons can also be described as "reflected" from the diffusive layer, often referred to as "diffuse reflectance" in other studies (*42*, *51*, *52*). This behavior allows us to apply Beer’s law to approximate tissue interactions and interpret our measurements as backscattering or diffuse reflectance originating from photons that were not absorbed within the diffusive layer. The diffusive layer can thus be treated as a homogeneous virtual light source within the tissue, denoted as *I*_0_. It is important to note that while UV light can penetrate deeper into the tissue, photons scattering into these deeper regions are more likely to be absorbed before returning to the surface and contributing to the detected signal. This provides tissue sectioning capabilities of the tissue surface layer.

Determining the exact distance between the diffusive layer and the tissue surface is challenging due to the limited availability of data on tissue properties in the deep UV range and the significant variability in reported values across studies (*50*, *53*, *54*). In this work, we employ oblique illumination at ∼70° to the optical axis to enhance optical sectioning in the UV range (*55*) and avoid detection of the reflected light from the quartz slide. Under this configuration, we assume the distance between the diffusive layer and the tissue surface is comparable to the depth of focus (DOF) of our objective (∼6 µm). This assumption is supported by the sharpness of our 255 nm and 280 nm images, which would appear blurry if the diffusive layer were significantly deeper than the DOF. Fig. S2 further demonstrates this relationship, showing that illumination closer to the optical axis (45° and 30° instead of 70°) results in diminished image sharpness, reinforcing the connection between the depth of the diffusive layer and the DOF. A more comprehensive discussion on modeling tissue behavior in the deep UV region is provided in the Methods section and Supplementary Material (Text S1).

For molecular quantification, we adopt the assumptions of Zeskind, et al., (*41*) that nucleic acids and proteins are the dominant absorbing biomolecules at both 255 nm and 280 nm, and we therefore include only their concentrations (and, by extension, masses) in the Beer’s law model. For our imaging system, *I*_0_ is calibrated against known average nuclear concentrations of nucleic acids and proteins, as well as polystyrene beads (see Materials and Methods). The calibrated *I*_0_ is then used to compute nucleic acid and protein mass across each absorption image via Beer’s law. The tryptophan quantum yield map (Fig. 1J) is derived from the estimated protein concentration in conjunction with the tryptophan autofluorescence image (see Materials and Methods).

### Label-free epi-DUV absorption and autofluorescence imaging of fresh mouse brain

Figure 2 presents images of a fresh mouse brain acquired with the three independent channels: 255A (Fig. 2A), 280A (Fig. 2B), and tryptophan autofluorescence at 357 nm (357F) (Fig. 2C). The top row shows whole-brain views with colored boxes indicating three regions of interest. Subsequent rows correspond to zoomed-in views of the stratum pyramidale of the hippocampus (Fig. 2D), lateral thalamus (Fig. 2E), and pretectal thalamus (Fig. 2F), based on corresponding regions in the whole mouse brain atlas (*56*). Across all regions, the 255A channel provides the strongest nuclear contrast and reveals very fine structural detail, consistent with the absorption peak of nucleic acids near 255 nm. For example, densely packed cells surrounding the hippocampus with clear nuclear contrast (red arrow) and the structure of a more myelinated region (black arrow) are clearly visible. The 280A channel shows slightly reduced nuclear contrast but emphasizes protein absorption. This difference is particularly evident in the pretectal thalamus (Fig. 2F), where some of the darker, more absorptive areas in 280A deviate from those observed in 255A. This can be attributed to variations in protein distribution across different regions of the mouse brain, such as dopamine receptor proteins (D1), which are highly concentrated in the thalamus but not the hippocampus (*57*). The 357F channel provides the least nuclear contrast but highlights regions with strong tryptophan autofluorescence, reflecting high local tryptophan concentrations, such as in the hypothalamus (*58*). The reduced nuclei contrast in the autofluorescence images is likely caused by a mismatch between the UV sectioning thickness and the DOF of the objective. Specifically, while DUV light that penetrates beyond the DUV “diffusive” layer is nearly completely attenuated, autofluorescence generated at these deeper regions will not, thus generating more out-of-focus signal. Therefore, the epi-DUV images do have better sectioning capabilities than autofluorescence (and also fluorescence if the tissue were labeled). Further details are provided in the Materials and Methods section.

**Fig. 2.**
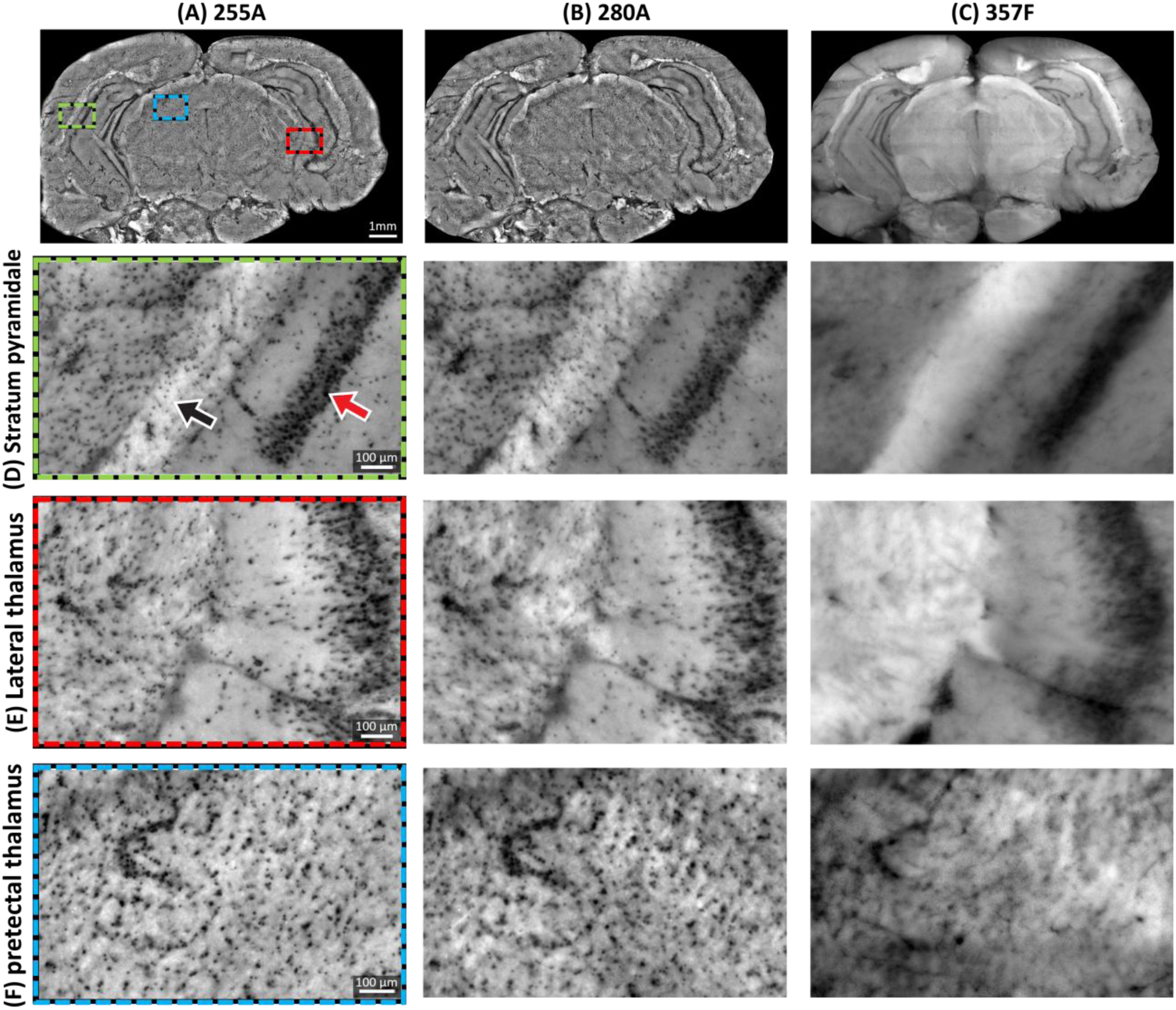
Label-free epi-DUV absorption and tryptophan autofluorescence images from fresh mouse brain. (**A**) Column for 255A images. (**B**) Column for 280A images. (**C**) Column for 357F images. (**D**) Row for zoomed-in region representing the stratum pyramidale of the hippocampus. The red arrow corresponds to densely packed cells surrounding the hippocampus. The black arrow corresponds to a more myelinated region. (**E**) Row for zoomed-in region representing the lateral thalamus. (**F**) Row for zoomed-in region representing the pretectal thalamus.

### Quantitative maps of representative regions in the fresh mouse brain

The quantitative maps are generated from the label-free epi-DUV absorption images and the autofluorescence images using Beer’s law. Figure 3 presents quantitative maps for the zoomed-in regions from Fig. 2 in the same order. Specifically, Fig. 3A, 3B and 3C respectively show the nucleic acid mass, protein mass, and quantum yield of tryptophan for the stratum pyramidale of hippocampus (Fig. 3D), lateral thalamus (Fig. 3E), and pretectal thalamus (Fig. 3F). The nucleic acid and protein mass maps reveal nuclei and protein-rich domains, respectively. Notably, the protein mass map also highlights certain nuclei, reflecting the protein content within nuclei, but with lower clarity than the nucleic acid map, as the protein map emphasizes overall protein distribution across the tissue. Quantum yield map highlights areas with high tryptophan autofluorescence efficiency. The resulting quantitative values are consistent with prior reports from UV transmission imaging on live cells (*40*, *41*). Specifically, the nucleic acid mass and protein mass, measured here, are up to 30 fg and 450 fg, respectively, per projected pixel in the object space (0.062 µm^2^) and a quantum yield of 15%. When scaled to the pixel size used in this study (see Materials and Methods), these values are in excellent agreement with those reported in the literature for thin samples (*40*, *41*), which show a range of max mass values of 17-50 fg for nucleic acid, 170 - 620 fg for protein, and up to 20% quantum yield. This validates the assumptions of modeling tissue behavior using Beer’s law within the diffusive layer near the tissue surface in absorption images and reveals the potential of molecular quantitative mass mapping of thick tissues.

**Fig. 3.**
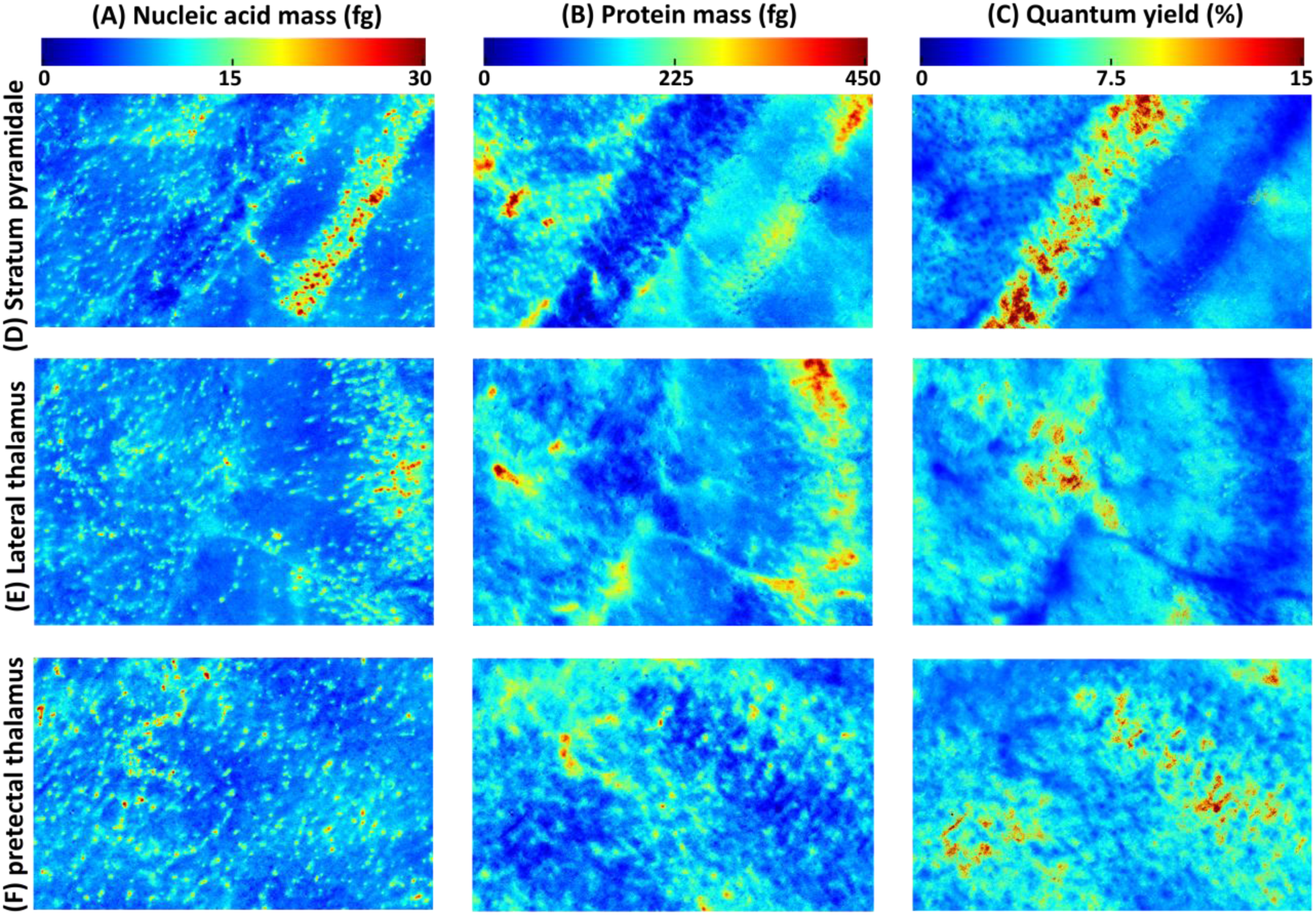
Quantitative maps of zoomed-in regions in Fig. 2 from fresh mouse brain sample 1. (**A**) Column for nucleic acid mass in femtograms. (**B**) Column for protein mass in femtograms. (**C**) Column for quantum yield of tryptophan in percentage. (**D**) Row for a region representing the stratum pyramidale of the hippocampus. (**E**) Row for a region representing the lateral thalamus. (**F**) Row for a region representing the pretectal thalamus.

### Pseudo-colorization via single-channel histogram matching

Beyond providing quantitative values from label-free epi-DUV images, we also apply a single-channel, fast transformation from the 255A image (Fig. 4A) to a pseudo–H&E image (Fig. 4B) that is familiar to pathologists. The method uses an efficient and simple histogram-matching transformation to align the 255A image to each RGB channel of a reference H&E image (Fig. 4C), then concatenates the three matched color channel images (all derived from 255A images only) to form the pseudo–H&E result. Prior work using dark-field reflectance ultraviolet microscopy (DRUM) (*42*) also produced pseudo-colorized images by combining 255A and 357F data with optical-density coefficients of H&E derived from Ruifrok (*59*) (Fig. 4D). Compared with the DRUM approach, our method offers: (1) greater speed and simplicity by requiring only the 255A channel (no autofluorescence); (2) higher nuclear contrast and improved color rendition; and (3) flexibility to match any desired H&E color palette without using fixed transformation coefficients. Figures 4A-D show representative figures of each image mode, with Figs. 4E-H illustrating one zoomed-in hippocampus region. Note the similar color tone between our pseudo-H&E image derived from the 255A image alone (Fig. 4B and 4F) and the real H&E image (Fig. 4C and 4G), both with clear nuclear contrast. Pseudo-H&E images derived from 255A and 357F (Fig. 4D and 4H) exhibit a large color tone mismatch and blurrier background, due to the use of the fluorescence image as a counterstain. Additional images showing a myelinated region (Fig. 4I–L) and posterior thalamus region (Fig. 4M–P) further support the superior color transformation from our single-channel 255A images.

**Fig. 4.**
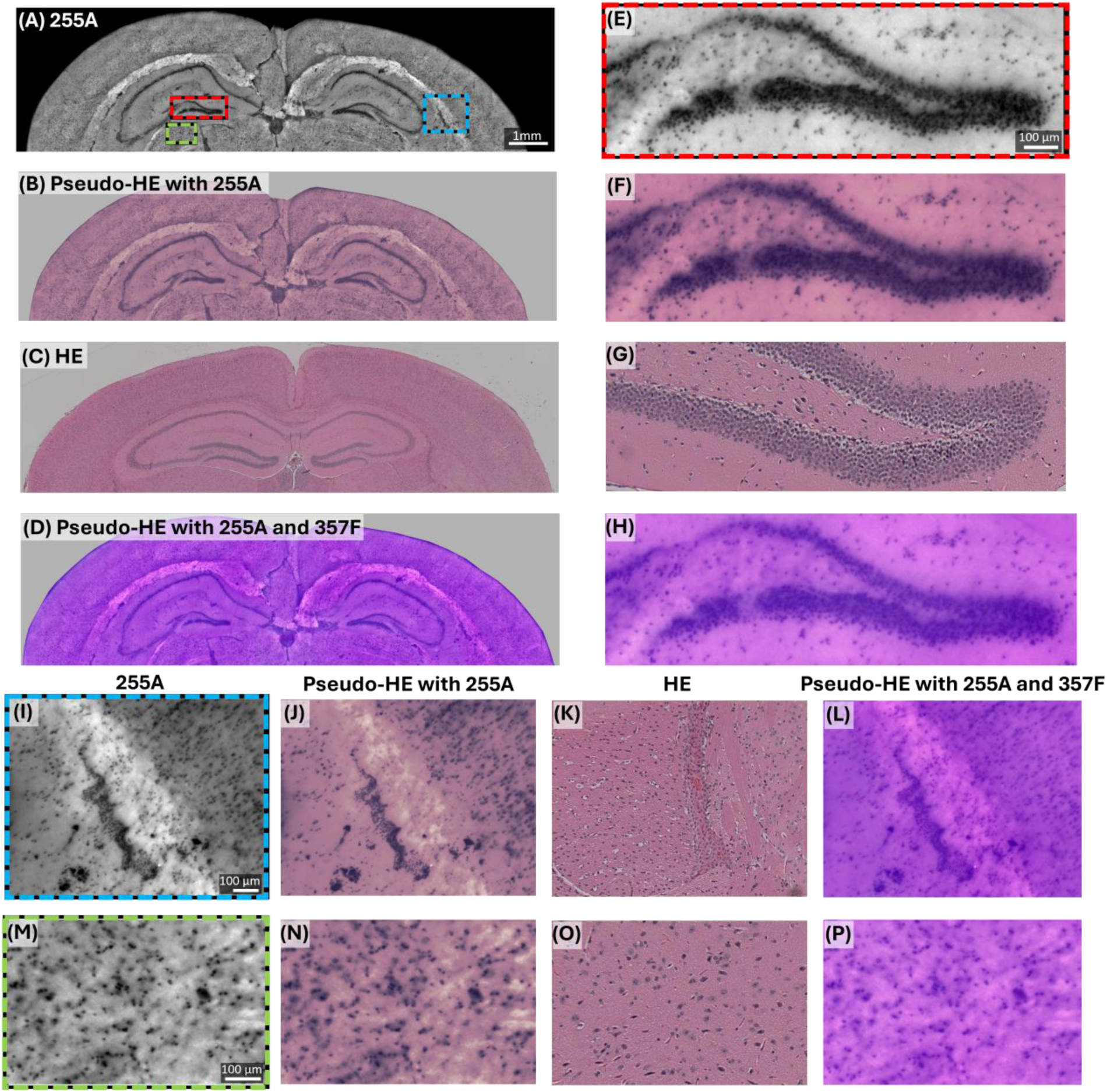
Pseudo-H&E colorization using single channel 255A transformation and comparison to DRUM Pseudo-H&E colorization. (**A**) 255A image of the mouse brain. (**B**) Pseudo-H&E image generated with 255A. (**C**) Brightfield image of an adjacent H&E-stained thin tissue section. (**D**) Pseudo-H&E image generated with both 255A and 357F (DRUM method). (**E**) Zoomed-in regions of 255A showing nuclei surrounding the hippocampus. (**F–H**) Pseudo-H&E transformed with 255A, adjacent H&E-stained thin tissue section, and DRUM pseudo-H&E of the hippocampus region, respectively. (**I**) 255A of another zoomed-in region from a myelinated area. (**J–L**) Pseudo-H&E transformed with 255A, adjacent H&E-stained thin tissue section, and DRUM pseudo-H&E of the myelinated area, respectively. (**M**) 255A of another zoomed-in region corresponding to the posterior thalamus. (**N–P**) Pseudo-H&E transformed with 255A, adjacent H&E-stained thin tissue section, and DRUM pseudo-H&E of the posterior thalamus region, respectively.

### Label-free epi-DUV images, quantitative maps, and pseudo-H&E of mouse kidney and human brain tumor samples

Beyond imaging the mouse brain, we apply our approach to mouse kidney and human brain tumor samples. The 255A (Fig. 5A) and 280A images of a cross-section of the surface of a whole mouse kidney demonstrate high nuclear contrast within the kidney tissue, with the pseudo-H&E image (Fig. 5B) closely resembling the adjacent H&E-stained slice (Fig. 5C). Notably, multiple glomeruli (black arrows) are distinctly visible in both the 255A and pseudo-H&E images (Fig. 5D), which are further confirmed by the adjacent H&E-stained thin slice of the same region (Fig. 5E), also showing clear glomerular structures (black arrows). Likewise, in the medulla region of the kidney, both the 255A and pseudo-H&E images (Fig. 5F, G) exhibit pronounced nuclear contrast, which is validated by an adjacent H&E-stained image of the same region (Fig. 5H). A zoomed-in view of a single glomerulus further highlights the nuclei contrast in the 255A (Fig. 5I) and pseudo-H&E images (Fig. 5J), which is again corroborated by the adjacent H&E-stained image of the glomerulus (Fig. 5K). Interestingly, the nucleic acid map (Fig. 5L) does not delineate the structure of the glomerulus as effectively as the protein mass map (Fig. 5M) and the quantum yield map (Fig. 5N), Notably, we find that the maximum protein mass is higher in kidney than in mouse brain; and, similarly, that the average protein mass in kidney is higher than in the brain, which is consistent with prior reports (*60*). This is further supported by comparing the average protein mass per unit area for mouse brain and mouse kidney samples, with results reported in the supplementary material (Fig. S5).

**Fig. 5.**
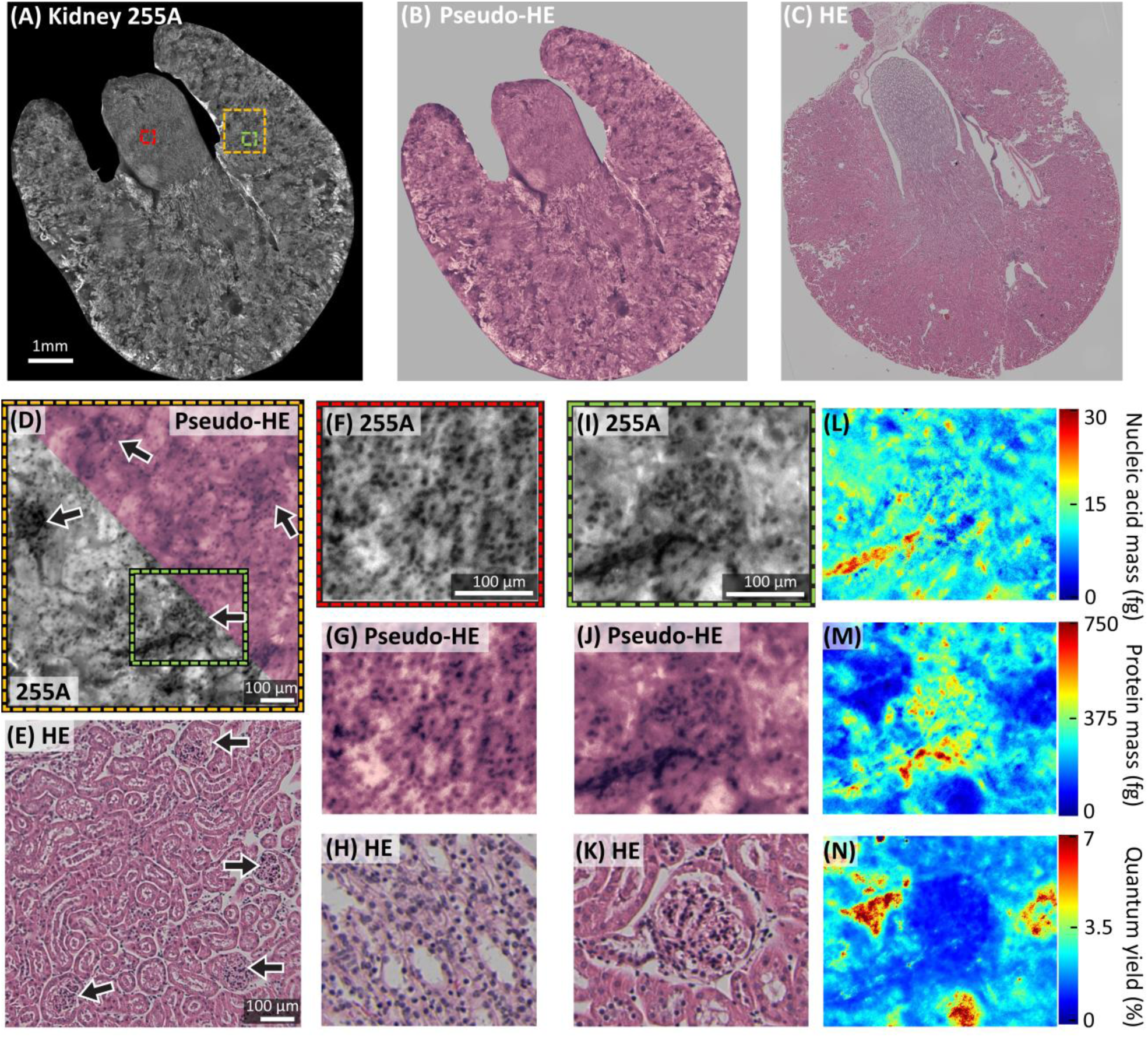
Epi-DUV images, pseudo-colorization, and quantitative maps of a mouse kidney sample. (**A**) 255A image of the whole mouse kidney sample. (**B**) Pseudo-H&E image of the kidney sample. (**C**) Adjacent H&E slice image of the kidney sample. (**D**) A zoomed-in region of (A), displaying multiple glomeruli (black arrows) in half 255A and half pseudo-H&E. (**E**) Adjacent H&E slice image with the same area, showing glomeruli (black arrows). (**F**) 255A image of another zoomed-in region of (A), representing the medulla of the kidney. (**G, H**) Pseudo-H&E and an adjacent H&E slice of this medulla region, respectively. (**I**) A zoomed-in region of (D), isolating a single glomerulus. (**J, K**) Pseudo-H&E image of the single glomerulus and an adjacent H&E slice of a single glomerulus. (**L–N**) Maps of nucleic acid mass, protein mass, and quantum yield of tryptophan of the single glomerulus image.

We also quantify the molecular mass maps of a human brain tumor sample (astrocytoma, grade 4). The images clearly exhibit densely packed cell nuclei, which aligns with diagnostic indicators of malignancy, such as uncontrolled cell proliferation and nuclear pleomorphism (Fig. 6A). A zoomed-in region (Fig. 6B) reveals clear nuclear contrast in the 255A image, the pseudo-H&E image, and nucleic acid mass map (Fig. 6C). A further zoomed-in region (Fig. 6D, E) highlights a dense tumor region, which is validated by the adjacent H&E-stained slice (Fig. 6F). Interestingly, the nucleic acid mass distribution (Fig. 6G) differs significantly from the protein mass distribution (Fig. 6H). This observation indicates that proteins in this malignant region are not predominantly localized within the cell nuclei, suggesting altered protein distributions in cancer (*61*). Additionally, this region exhibits both lower maximum and average quantum yield values (Fig. 6I) compared to the mouse brain, a difference that will be further analyzed in Fig. 7.

**Fig. 6.**
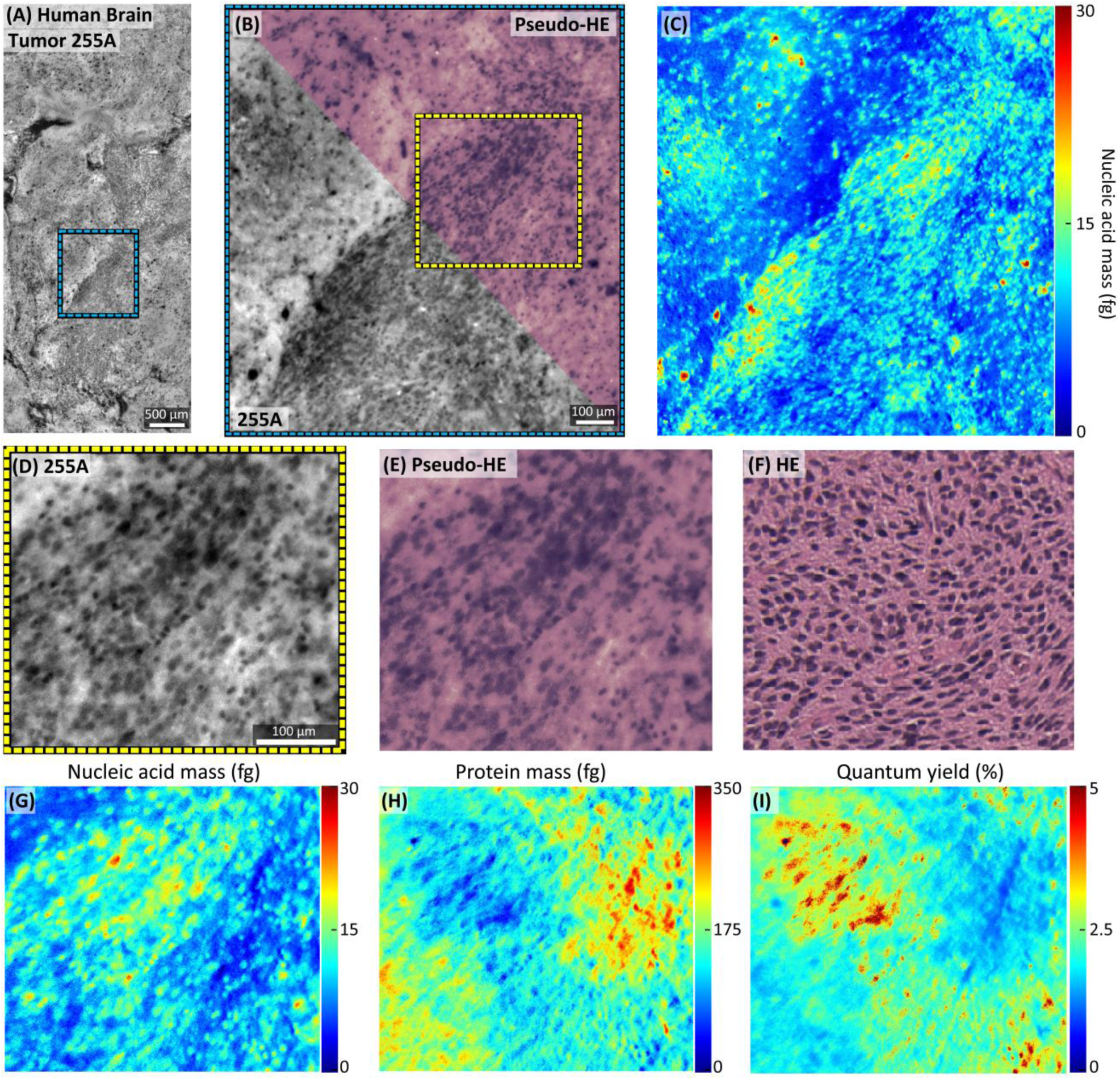
Epi-DUV images, pseudo-colorization, and quantitative maps of a human brain tumor sample. (**A**) 255A image of a brain tumor region. (**B**) A zoomed-in region corresponding to a dense tumor region. (**C**) The nucleic acid mass map of the region in (B), highlighting regions with high nuclear density and concentration of nucleic acid. (**D**) 255A image of a further zoomed-in region in (B). (**E**) Pseudo-H&E image of (D). (**F**) Adjacent H&E slice image of (D). (**G–I**) The nucleic acid mass map, protein mass map, and quantum yield of tryptophan of the region in (D).

**Fig. 7.**
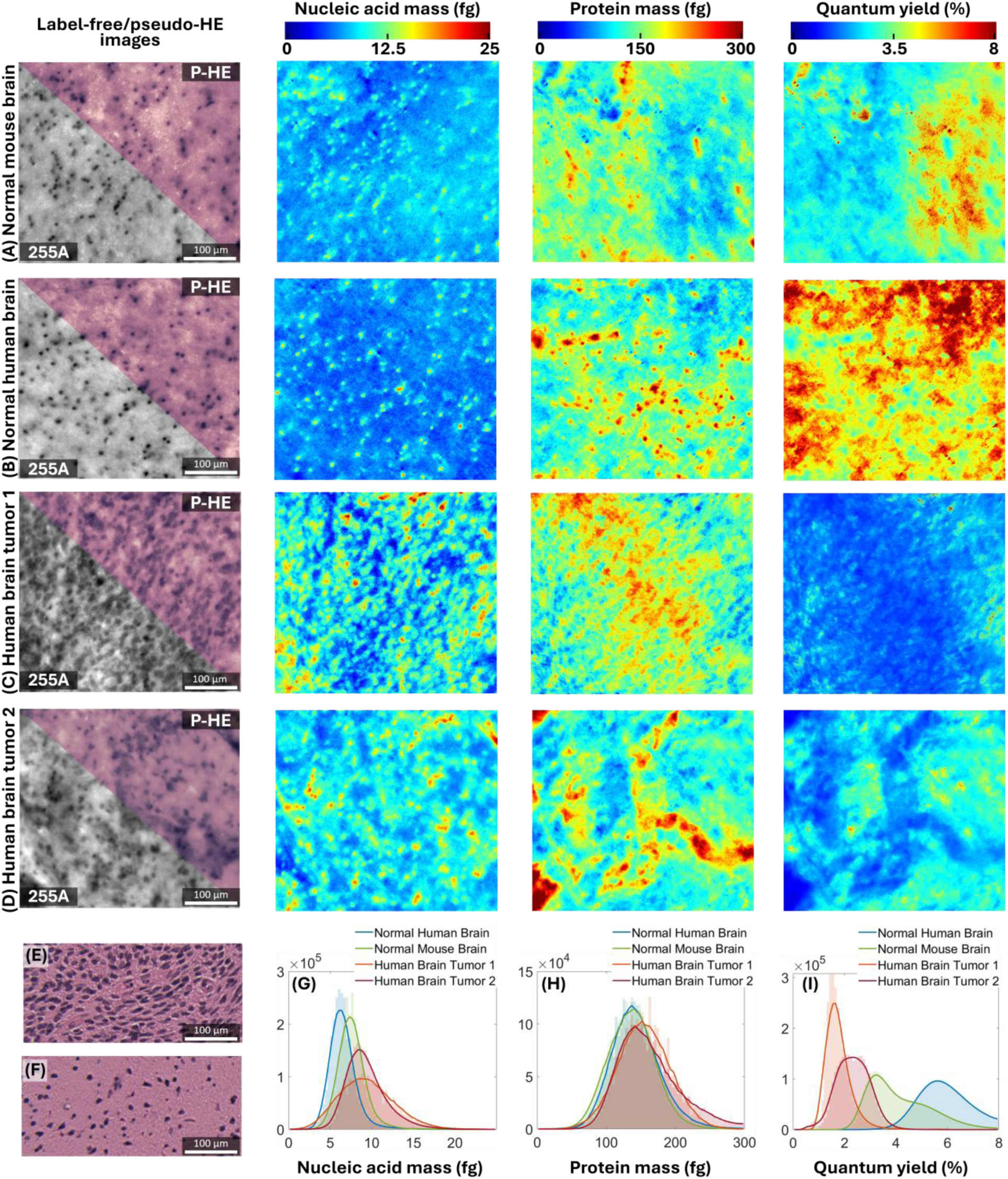
Comparison of quantitative values for brain tissues. (**A**) Row representing a cortex region in normal mouse brain, comprising (from left to right) 255A image with psuedo-H&E image (combined into one figure), nucleic acid mass, protein mass, and quantum yield. (**B**) Row representing a cortex region in normal human brain in the same sequence as (**A**). (**C**)-(**D**) Row representing cancerous region from two separate glioma samples in the same sequence as above. (**E**) Adjacent H&E slice image of the human brain tumor 1 sample. (**F**) Adjacent H&E slice image of the normal mouse brain cortex. (**G–I**) histogram of nucleic acid mass, protein mass, and quantum yield, respectively, of the 4 regions.

### Comparison of quantitative values for brain tissues

This section demonstrates how quantitative data can be utilized to compare brain tissues disease status. We present a comparison of a healthy mouse brain cortex region (Fig. 7A), a healthy human brain cortex region (Fig. 7B; adjacent tissue from a tumor excision), and tumor regions from two glioma patients (Figs. 7C, D). The cellularity of these images is corroborated by adjacent H&E images of a tumor region (Fig. 7E) and healthy mouse brain cortex (Fig. 7F).

The quantitative values reveal intriguing molecular differences between these tissues. For instance, the nucleic acid and protein mass distributions for the normal mouse and human brain cortex regions show notable similarities (see histograms, Figs. 7G, H). In contrast, the glioma regions (Figs. 7C, D) exhibit significantly higher cellularity than the healthy human brain cortex (Fig. 7B), as expected. Notably, the nuclei in the glioma tissues are not just more dense, they also display increased overall nucleic acid mass compared to the healthy tissues (see histogram, Fig. 7G). Interestingly, the protein content does not differ substantially between the healthy and cancerous tissues, whereas the quantum yield demonstrates a pronounced variation. The histogram of the quantum yield for the two cancerous regions and the healthy human brain (Fig. 7I) shows the average value in the human brain tumor is lower by more than half of the healthy tissue. This reduction may be attributed to differences in brain regions or alterations in the fluorescence properties of tryptophan due to the local tumor environment. However, these hypotheses remain speculative, as (to our knowledge) there are no existing literature characterizing changes in tryptophan quantum yield in tumor settings. This illustration, nevertheless, underscores how quantitative differences can be applied to gain new insight into the molecular composition of intact tissues and disease status.

## Discussion

In this study, we utilized epi-DUV microscopy to achieve label-free and slide-free histological imaging with quantitative mass mapping of nucleic acids and proteins, and tryptophan autofluorescence quantum yield using a multispectral approach. To the best of our knowledge, this approach represents the first demonstration of quantitative deep-UV molecular imaging in arbitrarily thick tissue samples using a simple widefield configuration. Combined with an extremely simple and rapid pseudo-H&E transformation using a single 255A image, our method offers the potential for real-time, quantitative evaluation of freshly resected or biopsy tissues. This capability could serve as an alternative to traditional histological workflows, providing clinicians with enhanced molecular information at the subcellular level for improved diagnostic and therapeutic decision-making.

Recent advancements have also demonstrated label- and slide-free histological imaging with virtual H&E staining on murine and human tissue samples using techniques such as reflectance confocal microscopy (RCM) (*62*), computational high-throughput autofluorescence microscopy with patterned illumination (CHAMP) (*63*), photoacoustic remote sensing (*26*, *27*), and photon absorption remote sensing (*64*, *65*). While promising, these methods rely on expensive laser systems, which significantly increase the cost and complexity of the systems. Additionally, these methods utilize neural networks for virtual staining, which can be time-consuming, require substantial amounts of training data, and are prone to hallucination errors, particularly when using unsupervised methods like CycleGAN (*66*). Moreover, the images generated from most of these techniques are generally not quantitative, which introduces additional issues when using virtual image translation techniques.

Quantitative oblique back-illumination microscopy (qOBM) has emerged as a powerful tool for label- and slide-free histological imaging (*67*) due to its ability to provide clear cellular and subcellular quantitative phase contrast of thick tissues in 3D using simple and inexpensive instrumentation (*68–72*). H&E-like image formation with qOBM, however, also relies on unsupervised neural networks, although it does have the advantage of enabling 3D H&E-like imaging, potentially in vivo (*67*).

As described in the Results, DRUM (*42*) employs diffusely reflected UV light at the tissue surface for qualitative imaging, and also enables label- and slide-free pseudo-H&E colorization that does not require neural networks. While similar, our approach offers several key advantages: (1) Pseudo-H&E transformation is achieved with a single 255A acquisition instead of multiple captures with different detection wavelengths. Our approach is not only faster and simpler, but it also yields better color matching to H&E and has improved nuclear contrast. (2) Our implementation of light pipes for illumination enables the use of low-cost UV objectives, as opposed to expensive long working distance objectives. (3) Most critically, our approach yields quantitative maps of nucleic acid mass, protein mass, and quantum yield, thus providing unique quantitative molecular information of intact tissues.

For quantification, our approach assumes that nucleic acids and proteins are the only two biomolecules absorbing at 255 nm and 280 nm. This assumption is reasonable since these biomolecules are indeed the dominant absorbers at these wavelengths, as has been reported in previous studies (*40*, *41*). Nevertheless, in complex tissue such as those analyzed in this work, this simplification occasionally results in negative mass values of both the nucleic acid and protein mass maps. These negative values suggest the presence of other strong UV absorbers, such as lipids, that were not accounted for in our model. However, the negative values observed here represent less than 1% of the total pixels within a given field of view (FOV). For better visualization and adherence to our assumption, we set all negative mass values to zero in the mass maps.

To improve the accuracy of the mass model, additional wavelengths could be introduced, such as 239 nm, which corresponds to the absorption peak of lipids (*73*). By experimentally determining the molar extinction coefficients of lipids at 239 nm, 255 nm, and 280 nm, lipid absorption in tissue could also be assessed. Similarly, additional wavelengths could be added to achieve a more faithful representation of the molecular composition of intact tissue.

The ability to obtain pseudo-H&E images of thick tissues using a single widefield acquisition via a simple RGB histogram equalization is significant and can have important implications for improving pathological workflows. Indeed, these maps (effectively of 255 nm absorption images of unprocessed tissues) are not identical to FFPE-processed H&E sections, but they recapitulate critical histological detail in a facile and fast manner. Results could be further enhanced by leveraging unsupervised neural networks. As discussed above, such methods do run the risk of hallucinations, but given the close similarity between our epi-DUV pseudo-H&E images and FFPE H&E sections, the likelihood of generating severe hallucinations is significantly lower. Plus, meaningful translation mistakes with epi-DUV to vH&E would be easy to detect, which is not the case with other label-free imaging modalities that rely more heavily on AI for image translation.

Finally, as we have explored in previous work (*34*, *35*), we highlight that single color (255A) DUV imaging of fixed tissue sections does not yield a realistic pseudo-H&E image, unlike fresh, unaltered tissues, indicating significant molecular compositional differences between fresh, unaltered tissues and fixed tissue sections. Such molecular changes have also been observer under phase imaging where the refractive index composition of FFPE processed tissues are significantly different than those of fresh tissues (*74*). More recently, using mid-infrared spectrochemical imaging (MIRSI) (*75*),it was found that FFPE tissues have substantially altered chemical/molecular and morphological compositional differences compared to frozen sections and—in particular—fresh tissues. Together, these findings underscore the importance of methods that can provide label-free structural and molecular information of intact/unaltered (fresh, unstained) tissues. The results of this work show epi-DUV microscopy is ideally suited to address these challenges.

In summary, we have developed a novel, LED-based epi-DUV microscopy system that enables rapid, label-free, and slide-free quantitative imaging of fresh, thick tissue samples. By leveraging the intrinsic absorption and autofluorescence properties of biomolecules in the deep-UV spectrum, we achieve subcellular resolution and generate quantitative mass maps of nucleic acids, proteins, and tryptophan quantum yield, alongside realistic pseudo-H&E images. This work highlights the potential of epi-DUV microscopy as a transformative tool in histopathology. Not only does this approach bypass the laborious histological workflow associated with FFPE and H&E staining, it also provides additional quantitative molecular information that could enhance diagnostic accuracy and therapeutic decision-making. Ultimately, this versatile and accessible imaging system has great potential to revolutionize clinical pathology, providing holistic, real-time evaluation of fresh tissues while also remaining simple (widefield), label-free and low cost, maintaining compatibility with low-resource settings.

## Materials and Methods

### Label-free epi-DUV imaging system

The epi-DUV system is built on a conventional wide-field inverted microscope setup (Fig. 1A) with a modified illumination path incorporating LEDs and light guides (Fig. 1B). Narrow-band surface mount LEDs with center wavelengths of 255 nm (Photon Wave Inc.) and 280 nm (Nichia Inc.) were used. These LEDs were mounted on heat sinks and secured below the sample stage using customized 3D-printed LED holders (Fig. 1B). Light pipes (63-093, Edmund Inc.) with a 4 mm aperture were inserted into the LED holders to guide the light from the LEDs, ensuring uniform illumination at the sample. The light pipes are oriented at a 70° angle to the optical axis, effectively operating in a dark-field configuration, as specular reflections fall outside the NA of the objective. The power of the two 255 nm LEDs and two 280 nm LEDs at the sample plane was controlled to be approximately equal (∼7.1 mW). The system lateral resolutions at 255 nm and 280 nm were measured to be ∼550 nm and ∼600 nm, respectively, as detailed in the supplementary material (Fig. S1).

Light information from the sample was collected using a UV-transmissive objective (LMU-15X-UVB, NA = 0.31, Thorlabs Inc.), passed through band-pass filters centered at 255 nm, 280 nm, and 357 nm (FF01-254/8-25, BrightLine Inc.; FF01-280/10, BrightLine Inc.; 18-384, Edmund Inc.), which were installed on a filter slider (CFS1, Thorlabs Inc.), a UV-transmissive tube lens (f = 150 mm), and finally captured by a UV-sensitive scientific complementary metal-oxide-semiconductor (sCMOS) camera (ATX081S-UC, 2840 × 2840 pixels, 2.74-µm pixel size, Lucid Inc.).

### Sample collection, image acquisition, and image processing

This study examined three types of tissues: mouse brain, mouse kidney, and human brain tumor. Mouse brain and kidney samples were obtained in accordance with the Institutional Animal Care and Use Committee of the Georgia Institute of Technology. Human brain tumor samples were obtained from the Winship Cancer Institute of Emory University under approved, IRB exempt protocols. All tissues were de-identified and imaged fresh, typically within six hours of removal. The two human brain tumor samples were confirmed post-surgery as a grade 4 astrocytoma with isocitrate dehydrogenase (IDH) mutation, classified according to the World Health Organization Classification of Tumors of the Central Nervous System.

For image acquisition, fresh tissue samples were placed on a quartz slide and gently compressed with a quartz coverslip or another quartz slide (depending on the rigidity of the sample) to flatten the bottom surface, minimizing surface irregularities. The quartz slide was secured on a sample stage (MPRC & MPSH2, Thorlabs Inc.), connected to a 3-axis translation stage (LX30, Thorlabs Inc.). Tissue scanning was automated using stepper motors, motor drive shafts, and motor drivers controlled via a custom MATLAB application interfaced with a microcontroller (Arduino Mega 2560 REV3). The tissue was scanned with approximately 20% overlap between adjacent FOVs to facilitate image stitching.

Each FOV measured ∼707 µm × 707 µm, and the scanning time for a 10 mm × 10 mm tissue sample at an exposure time of ∼330 ms was ∼5 minutes per wavelength. This throughput is comparable to that of a commercial digital scanner. Here we have a 1-second interval between image acquisitions of adjacent FOVs to allow the motor to settle and minimize stage vibrations caused by motor movement. Future improvements in motor-stage synchronization could further increase scanning rates.

To minimize stitching artifacts caused by uneven illumination during image stitching, each individual FOV is normalized using a background image. This background image is created by applying a severe Gaussian filter (*σ* = 1000 pixels) to each FOV, thus retaining only very low spatial frequency information. Dividing each FOV by its Gaussian-filtered version produces a normalized FOV with a highly uniform illumination distribution. This process has been referred to as flat-field correction (*76*).

H&E images for all tissue types were acquired digitally using a commercial slide and plate scanner at 10× magnification (NA = 0.3) (BioTek Cytation 7, Agilent Technologies Inc.).

### Quantification of nucleic acid and protein masses

As briefly described in the Results section, the quantitative mass maps were computed based on Beer’s law, assuming a diffusive layer at a specific depth within the tissue acts as a virtual light source for UV illumination. This assumption, combined with the strong absorption characteristics of biological tissues in the deep-UV range, enables the behavior of light within the tissue to be modeled as predominantly absorptive within the diffusive layer. Consequently, the measurements in the 255A and 280A images primarily reflect photons that were not absorbed at these wavelengths, following principles similar to conventional transmission UV microscopy. Under this model, the absorbance at each wavelength can be expressed using Beer’s law (*41*), assuming that nucleic acids and proteins are the only biomolecules contributing to absorption at 255 nm and 280 nm:

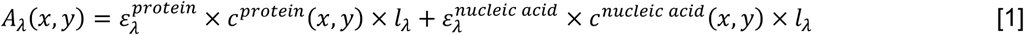

where *A*_*λ*_ (*x*, *y*) is the wavelength-dependent absorbance at each pixel, 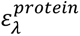 is the wavelength-dependent molar extinction coefficient of protein, *c*^*protein*^(*x*, *y*) is the concentration of protein at each pixel, *l*_*λ*_ is the wavelength-dependent path length, 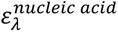 is the wavelength-dependent molar extinction coefficient of nucleic acid, and *c*^*nucleic*^ ^*acid*^(*x*, *y*) is the concentration of nucleic acid at each pixel. The path length is estimated to be approximately equivalent to the DOF of the objective, calculated to be ∼6 µm (*77*). As shown in the Supplemental Material (Text S1), the UV penetration depth of diffusely reflected light can reasonably be assumed to be comparable to the DOF. If this assumption were incorrect, there would be significant out-of-focus information within the FOV, which is not observed in the sharp, high contrast 255A and 280A images, where nuclei are clearly visible. (In fact, with less oblique illumination, for example, using 45° or 30° instead of 70°, appreciably more out-of-focus background light is indeed observed—see Fig. S2.) Additionally, the relationship between penetration depth and DOF with respect to the focus quality of the image has been discussed in prior studies (*63*).

Absorbance can be rewritten as:

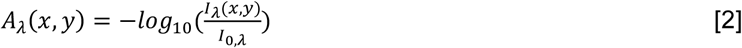

Where 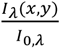 is the wavelength-dependent transmittance at each pixel. *I*_*λ*_ (*x*, *y*) is the wavelength-dependent transmitted intensity at each pixel, representing either the 255A or 280A images, and *I*_0,*λ*_ is the wavelength-dependent intensity at the diffusive layer at each pixel (i.e., the intensity of the virtual light source). To find the *pixel-wise* concentration of protein and nucleic acid, we must first calibrate *I*_0,*λ*_ of the epi-DUV system. It can be computed by assuming known *average* concentrations of protein and nucleic acid in cell nuclei, which are estimated from the literature to be ∼300 mg/ml and ∼30 mg/ml (*78*), respectively. The concentrations can be converted into molarity (M) based on the average molar mass of protein (52728 Da) and nucleic acid (330 Da) (*41*). With this, Eq [1] can be rearranged to estimate *I*_0,*λ*_:

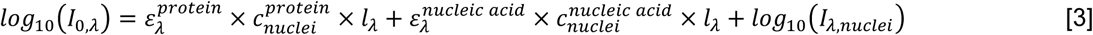

*I*_255,*nuclei*_, *I*_280,*nuclei*_ represent the *average* values of over 50,000 segmented visible nuclei in the stitched 255A and 280A images of a mouse brain sample. Using these values and the quantities described in Table 1, *I*_0_ at 255 nm and 280 nm can be calculated. In this study, *I*_0_ is assumed to be constant for all tissues analyzed, as the absorption properties of these biological tissues are comparable (*50*). Based on this assumption, we applied the calculated *I*_0_ to independent mouse brain samples, mouse kidney, and human brain tumor tissues. We also repeated the procedure by calculating the quantitative values for the mouse kidney using the *I*_0_ derived from independent kidney tissue, which produced results very similar to those obtained using the *I*_0_ calculated from the mouse brain, as shown in the supplementary material (Fig. S4).

**Table 1.** Quantities for computing mass maps and autofluorescence quantum yield of tryptophan. All molar extinction coefficients, *N*_*aa*_, and *F*_*Trp*_ were referenced from (*41*).

| Quantities | Values | Units |
| --- | --- | --- |
| $\epsilon_{255}^{protein}$ | 36057 | $M^{-1}cm^{-1}$ |
| $\epsilon_{280}^{protein}$ | 54129 | $M^{-1}cm^{-1}$ |
| $\epsilon_{255}^{nucleic\ acid}$ | 7000 | $M^{-1}cm^{-1}$ |
| $\epsilon_{280}^{nucleic\ acid}$ | 3500 | $M^{-1}cm^{-1}$ |
| $\epsilon_{255}^{Trp}$ | 3787 | $M^{-1}cm^{-1}$ |
| $c_{nuclei}^{protein}$ | 0.0057 | $M$ |
| $c_{nuclei}^{nucleic\ acid}$ | 0.091 | $M$ |
| $l_{255}$ | 5.84 | $\mu m$ |
| $l_{280}$ | 6.41 | $\mu m$ |
| $T_{AF}$ | 330 | $ms$ |
| $T_{trans}$ | 10 | $ms$ |
| $TE_{AF}$ | 0.75 | |
| $TE_{trans}$ | 0.48 | |
| $NA$ | 0.31 | |
| $N_{aa}$ | 466 | |
| $F_{Trp}$ | 0.014 | |

To further validate the calibrated *I*_0_ from the mouse brain, we used 10 µm polystyrene beads. First we measured the absorbance of the beads in transmission mode. Then, we placed the beads underneath fresh brain tissue, and using the same procedure described above, we computed *I*_0_ for our “epi mode” approach using the beads as reference (instead of cell nuclei as described above). We performed this bead validation only at 255 nm, as the beads absorb significantly less at 280 nm (*79*), which could lead to inaccurate results. A statistical comparison between the *I*_0_ values calculated from polystyrene beads and the nuclei in mouse brain in individual FOVs is presented in the supplemental material (Fig. S3), and revealed no statistically significant differences.

With the calibrated *I*_0_ at 255 nm and 280 nm, we then calculated the concentrations of protein and nucleic acid *at each pixel* (in molarity, M) by solving Eq [1] at each wavelength. Multiplying these concentrations by the magnified pixel area (∼0.062 µm^2^) and an effective path length (defined as the average DOF) provided the quantities in moles. From these values, the masses of protein and nucleic acid were determined based on their respective molar masses. These masses are defined as the mass contained within the volume represented by each pixel, projected through the DOF.

### Quantification of the autofluorescence quantum yield of tryptophan

The quantification process for the quantum yield of tryptophan in transmission UV microscopy has been thoroughly characterized in (*41*). In this study, we employed a similar approach, with minor modifications to account for the epi-illumination setup. The autofluorescence intensity at each pixel can be expressed below:

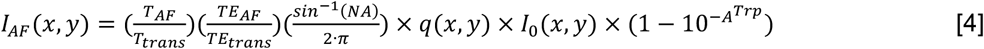

and

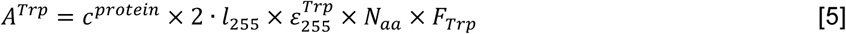

where *I*_*AF*_ (*x*, *y*) is the autofluorescence intensity of tryptophan at each pixel (357F images). *T*_*AF*_ and *T*_*trans*_ are the exposure times used in capturing the autofluorescence image and the transmission image, *TE*_*AF*_ and *TE*_*trans*_ are the transmission efficiency of the bandpass filters for autofluorescence and transmission at 357 nm and 255 nm, respectively, *NA* is the numerical aperture of the objective lens, *q*(*x*, *y*) is the autofluorescence quantum yield of tryptophan at each pixel, *I*_0_(*x*, *y*) represents the pixel values of the initial input light intensity at the sample plane within the FOV, *A*^*Trp*^ is the absorbance due to tryptophan, *c*^*protein*^ is the concentration of protein calculated for the mass map, *l*_255_ is the DOF of the objective at 255 nm (path length), 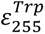 is the molar extinction coefficient of tryptophan at 255 nm, *N*_*aa*_ is the average number of amino acids per protein, and *F*_*Trp*_ is the average tryptophan frequency in amino acids. We estimated *I*_0_(*x*, *y*) by illuminating a blank quartz slide with a 255 nm LED in transmission mode, at the same power on the sample stage as in the epi-DUV setup (∼7.1 mW). This approach approximates the amount of light power entering the tissue for the 255 nm illumination in epi-mode. Note that this *I*_0_(*x*, *y*) differs from *I*_0,*λ*_, with the latter representing the intensity at the diffusive layer, which is used to compute mass maps. This distinction also deviates from the definition in (*41*) for transmission mode, where the same *I*_0_(*x*, *y*) was used to compute both the quantities of mass and quantum yield. The difference arises from the assumption that the initial light intensity is significantly attenuated before reaching the diffusive layer and that only a small portion of the scattered photons from the diffusive layer is collected, whereas nearly all the initial light contributes to potential tryptophan absorption and subsequent autofluorescence. This assumption is further supported by the fact that we used the same exposure time for both the absorption and autofluorescence images in epi-DUV, while in transmission-mode UV absorption microscopy, the exposure time is much shorter than that used for autofluorescence (*41*).

In the equation for *A*^*Trp*^, a factor of 2 is applied to the path length, reflecting the assumption that the 255 nm UV penetration depth that generates detected autofluorescence is approximately twice the depth for the diffusely backreflected 255 nm light that is detected in the absorption image (which is comparable to the DOF). This is because the latter undergoes a round trip, meaning it must travel back from the diffusive layer without being absorbed. Consequently, it is reasonable to assume that the light at the diffusive layer can also be scattered into deeper regions in the tissue at approximately the same distance. The exact penetration depth should ideally be modeled using more rigorous approaches such as MC simulations. However, estimating a UV penetration depth at approximately twice the DOF of our objective lens (∼12 µm) is a reasonable assumption, given the relatively extreme oblique illumination in our setup and previously reported data on UV penetration depths (*25*, *49*, *80*). All 255 nm light within this penetration depth can potentially be absorbed by tryptophan, thereby contributing to tryptophan autofluorescence. The mismatch between the UV sectioning thickness and the DOF is further supported by our 357F images, which appear noticeably blurrier compared to the 255A and 280A images.

With the above assumptions and the quantities described in Table 1, *q*(*x*, *y*) can be computed for each pixel according to equation [4].

### Comparison of quantitative values with previously reported data

In Fig. 3, our measured values for nucleic acid mass, protein mass, and quantum yield are up to 30 fg, 450 fg, and 15% per 0.062 µm^2^. This pixel size is 0.62 times the pixel size used in ref. (*40*) (0.1 µm^2^), where nucleic acid mass and protein mass of a live neutrophil were reported as up to 80 fg and 1000 fg, respectively. Adjusting for the pixel size difference, these values correspond to nucleic acid and protein masses of up to 50 fg and 620 fg, respectively, which align well with our results.

In another study (*41*), our pixel size is 1.72 times larger than their pixel size (0.036 µm^2^), where nucleic acid mass, protein mass, and the quantum yield of tryptophan for a live mouse macrophage cell were reported as up to 10 fg, 100 fg, and 12%, respectively. Scaling for the pixel size difference, these values correspond to nucleic acid mass, protein mass, and quantum yield of 17 fg, 170 fg, and 20%, respectively, which again align well with the range of our reported values.

Despite differences in cell types, our measured quantitative values show close agreement with those reported in the literature.

### Pseudo-colorization with 255A

The purpose of pseudo-colorization is to generate realistic H&E-like images with high nuclear contrast. The density and morphology of nuclei are among the most critical diagnostic features and are often sufficient for pathologists to make accurate diagnoses. The 255A images were selected for this transformation because they provide the highest nuclear contrast. A straightforward approach to performing the color transformation involves matching the histogram of the 255A image to that of a reference H&E image. This ensures that the dark nuclei in the 255A image correspond to the purplish nuclei in the H&E image, while the surrounding structures (that do not contain high nucleic acid concentrations) adopt a pinkish hue. This process can be efficiently executed using the built-in MATLAB function “imhistmatch”, which adjusts (shifts and stretches) the histogram of the 255A image to align with each of the RGB channels of the reference H&E image. The adjusted three channels were then concatenated to produce a pseudo-H&E image. This process is exceptionally simple, fast, and does not require post-processing, while successfully generating realistic H&E-like images with clear nuclei contrast. It is important to note that for consistent, high-quality histogram matching, the flat-field correction of the 255A image is critical, but the images do not need to be quantitative, i.e., normalizing by I_0_ is not necessary.

## Author contributions

Conceptualization: M.S., V.G., and F.E.R.

Methodology: M.S., V.G., A.S.T., A.R., B.E.H., and F.E.R.

Investigation: M.S., V.G., and F.E.R.

Visualization: M.S. and F.E.R.

Supervision: F.E.R.

Writing—original draft: M.S., V.G., and F.E.R.

Writing—review & editing: M.S. and F.E.R.

## Supplementary Information

**Supplementary text S1: Note on modelling absorption, scattering in the deep UV region, and epi-DUV effective sectioning**

We use Beer’s law to quantify the absorption images under the assumption that the “diffusive” layer (i.e., the virtual light source within the tissue) is homogeneous and immediately above the surface layer that is analyzed. We argue that these assumptions are reasonable, given that the acquired DUV absorption images do not suffer from significant out-of-focus contributions. We sought to further characterize UV penetration using Monte Carlo (MC) simulations; however, as highlighted in the Results section, the availability of reliable data on DUV scattering and absorption properties of intact tissues is limited and contains significant discrepancies across studies (*50*, *53*, *54*). One indicator of this unreliability is that reported values for absorption coefficients (∼11 − 13 *cm*^−1^) suggest that, in general, only one absorption event occurs every ∼770 − 910 *µm* in the deep UV range (*54*). This estimate is two orders of magnitude higher than the depth of focus of the objective used in this work (∼6 *μm*). If these values were accurate, our images would contain significant out-of-focus background even when using oblique or darkfield illumination (more out-of-focus contributions are observed when using *less* oblique illumination). Similar effects have also been observed using MUSE (*29*, *55*). This observation suggests that the absorption coefficient in the deep UV range is likely much higher than reported in the literature. While the exact optical properties of tissue in the deep UV range and the precise sectioning thickness remain uncertain, our absorption model is specifically focused on the region between the diffusive layer and the tissue surface. This layer is assumed to be very close to the surface under oblique illumination, with its distance comparable to the depth of focus (DOF) of the objective lens. Combined with the strong absorption behavior of deep UV light, this proximity allows us to approximate light behavior in this regime as predominantly absorptive and effectively modeled by Beer’s law.

Here, we provide an estimation of the absorption coefficient (*μ*_*a*_) of fresh mouse brain tissue at 255 nm. First, we calculate the theoretical *μ*_*a*_ of cell nuclei using Eq. 6:

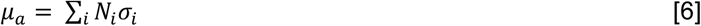

where *N*_*i*_ is the density of the *i-th* biomolecules in *cm*^−3^, which can be converted from known average concentrations of nucleic acid and protein in nuclei (∼30 *mg*/*ml*, ∼300 *mg*/*ml*, respectively (*78*)); *σ*_*i*_ is the absorption cross-section of the *i-th* biomolecules in *cm*^2^, which can be calculated from the molar extinction coefficient values of nucleic acid and protein at 255 nm. *μ*_*a*_ for nucleic acid is estimated to be ∼1466 *cm*^−1^ and that for protein is ∼472 *cm*^−1^. The sum of these values (∼1938 *cm*^−1^) is the total absorption coefficient in cell nuclei and should match the *μ*_*a*_ calculated from Beer’s law in Eq. 7 using the average segmented nuclei intensity and the calibrated *I*_0_. Note that *I*_0_ was estimated using (1) literature values of the concentration of nucleic acid and protein in nuclei (which should agree with the theoretical estimate of *μ*_*a*_ above, and (2) using polystyrene beads which makes no assumptions about the tissue. Both methods yielded the same value *I*_0_ (Fig. S3).

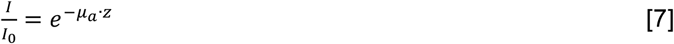

Where *I* is the *average* pixel intensity of segmented cell nuclei at 255 nm (over 50,000 nuclei were used here); *I*_0_ is the calibrated intensity at the “diffusive” layer (the virtual light source) at 255 nm; *z* is the path length (DOF of the objective lens) at 255 nm. *μ*_*a*_ is estimated to be ∼1941 *cm*^−1^ using these values, which agrees with the total *μ*_*a*_ of the nuclei computed from the absorption of nucleic acid and protein using Eq. 6.

We then substitute *I* in Eq. 7 with the average pixel intensity of a whole-brain 255A image of a fresh mouse brain, resulting in an estimation of *μ*_*a*_ = ∼1225 *cm*^−1^ for the mouse brain at 255 nm, which is two orders of magnitude higher than the value reported in (*54*). Using this value, the absorption length is estimated to be ∼8.16 *μm*. Considering the geometry of the tilted illumination source, the absorption length parallel to the optical axis is ∼6.23 *μm*, which agrees well with the DOF at 255 nm (∼5.84 *μm*).

***I***_**0**_. (**A**) Row of quantitative values of a region calculated using the mouse brain *I*_0_. (**B**) Row of quantitative values of the same region calculated using the mouse kidney *I*_0_. High agreement was observed in protein mass and quantum yield maps, while slight differences were present in the nucleic acid mass map.

**Fig. S1.**
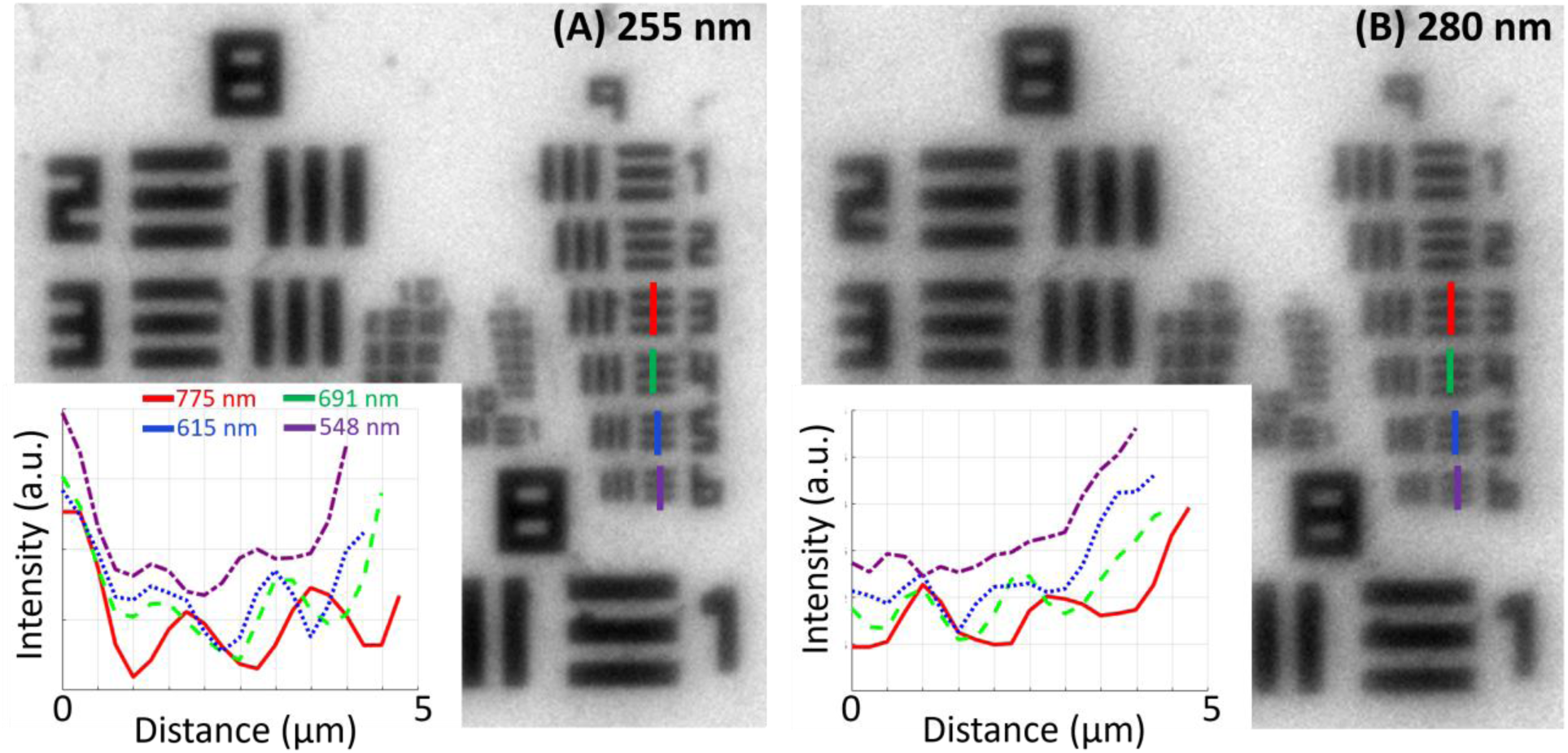
Experimental resolution testing using high-resolution USAF test target (Newport). (**A**) Resolution at 255 nm. (**B**) Resolution at 280 nm. Line plots corresponding to a resolution of 775 nm (red), 691 nm (green), 615 nm (blue), and 548 nm (purple). These measurements were performed in transmission mode.

**Fig. S2.**
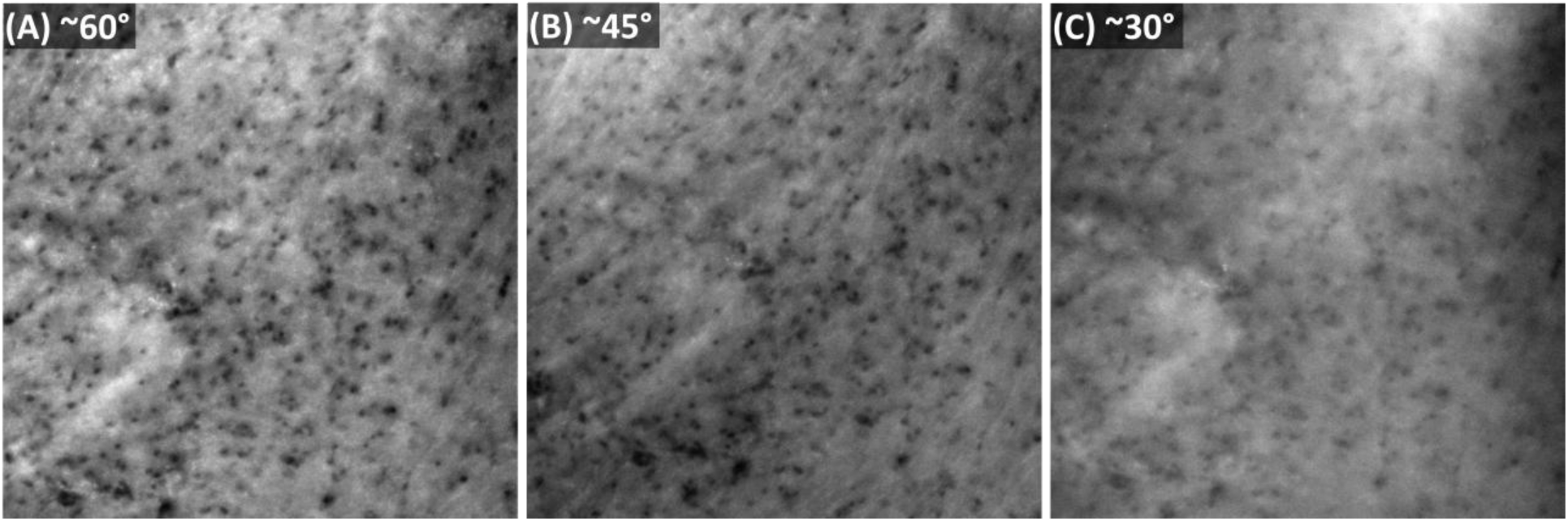
Illumination angle-dependent contrast in a 280A image. (**A**) Illumination with a light pipe at an angle of ∼60° to the optical axis. (**B**) Illumination with a light pipe at an angle of ∼45° to the optical axis. (**C**) Illumination with a light pipe at an angle of ∼30° to the optical axis.

**Fig. S3.**
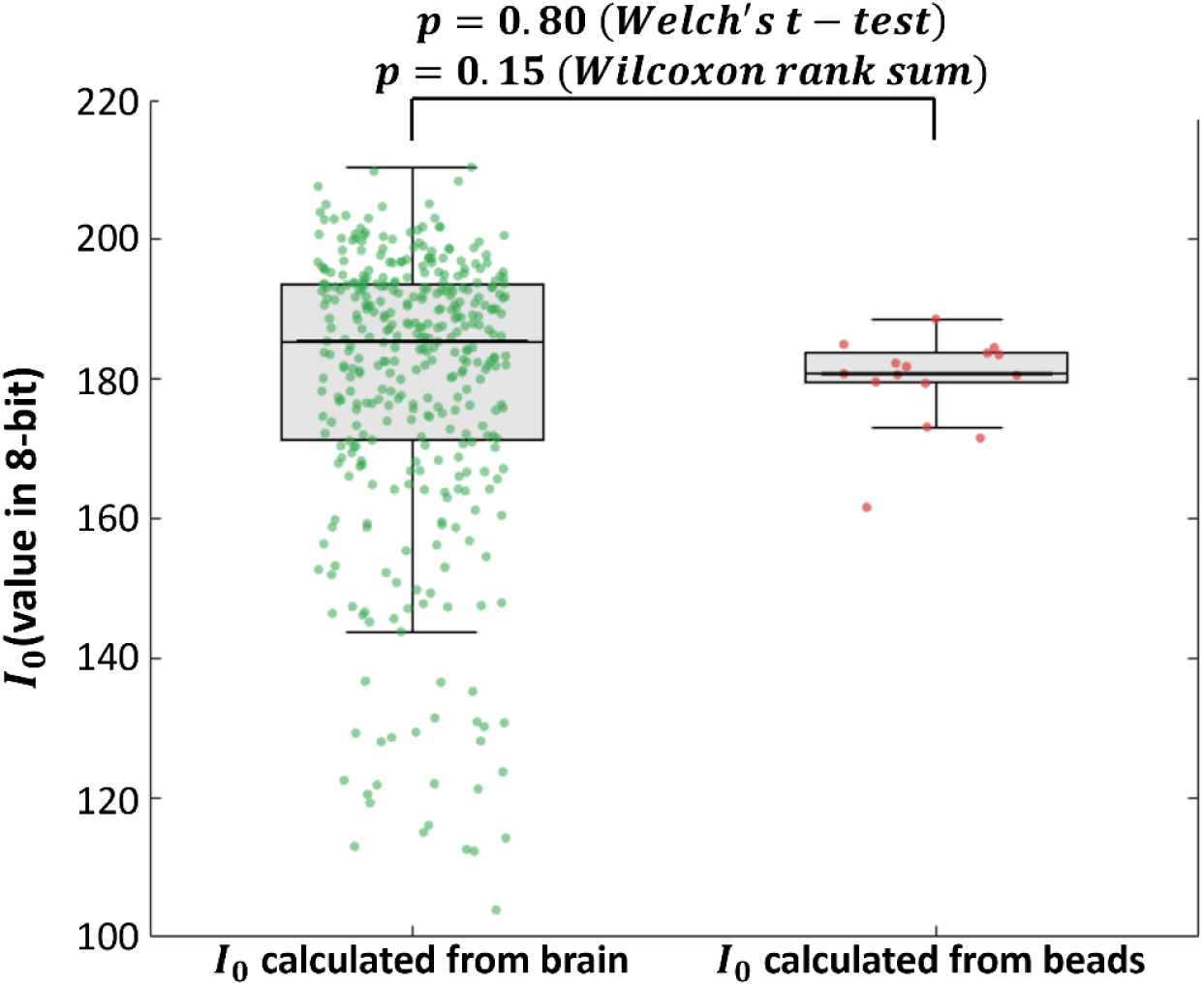
*I*_0_ validation for 255A by comparing distributions of *I*_0_ computed from the mouse brain and polystyrene beads. Welch’s t-test and Wilcoxon rank sum tests were conducted with n = 375 for the brain group and n = 15 for the bead group. No statistically significant differences were observed between the two groups (the significance is defined as *p* ≤ 0.05).

**Fig. S4.**
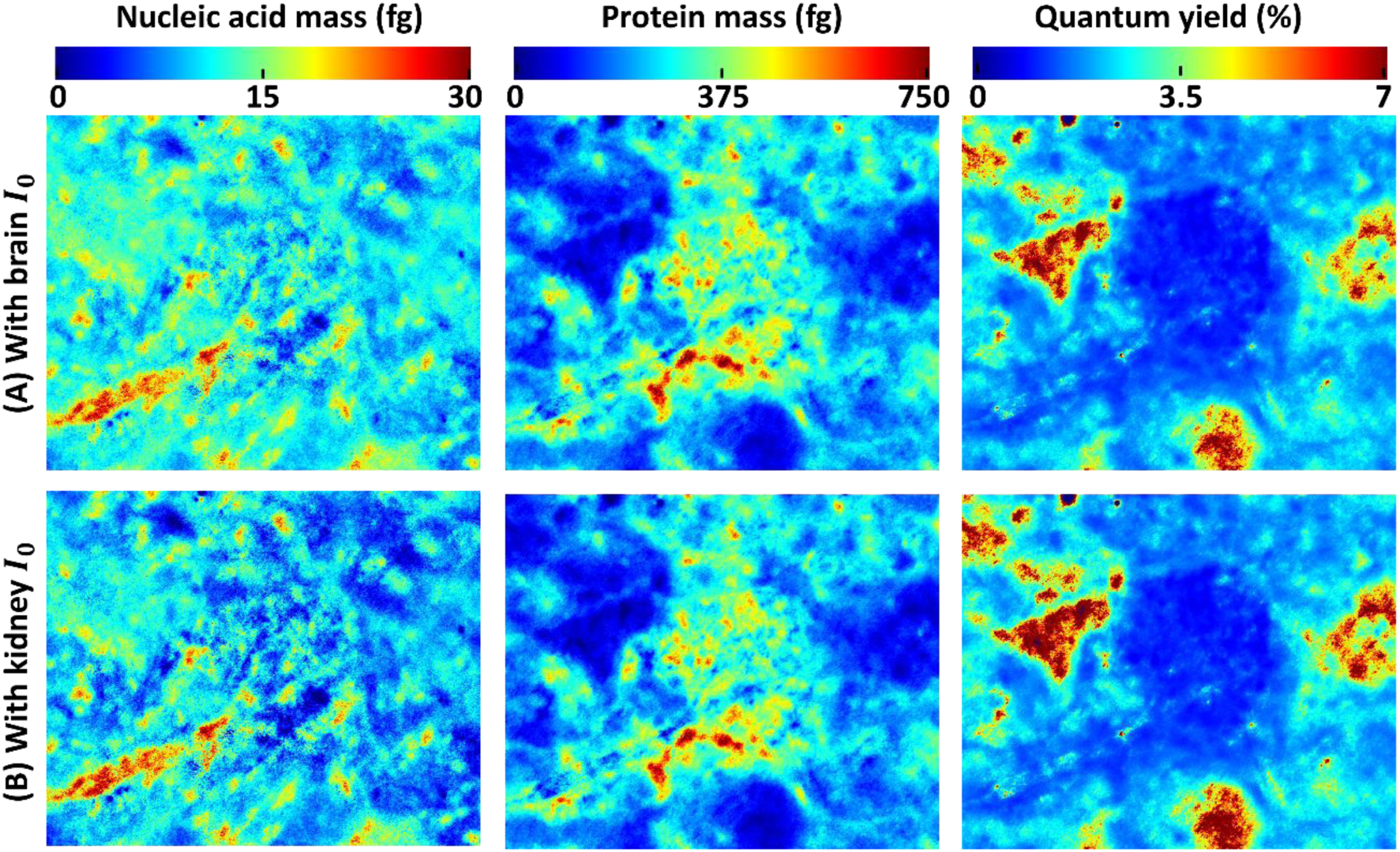
Comparison of quantitative values of a kidney region calculated using the mouse brain *I*_0_ and kidney.

**Fig. S5.**
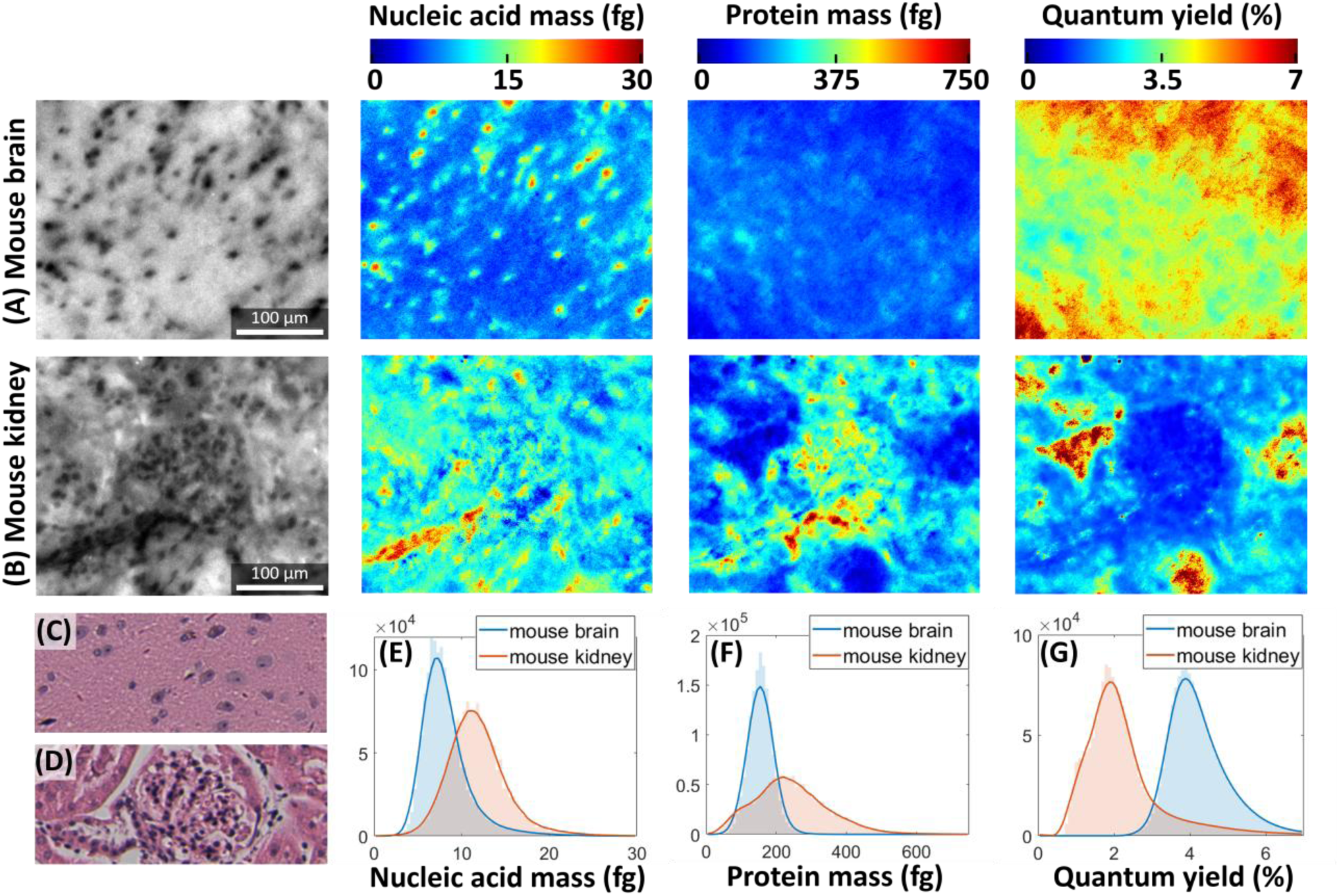
Comparison of quantitative values between the mouse brain and mouse kidney. (**A**) Row of a region in the mouse brain thalamus. (**B**) Row of a region in the mouse kidney, showing a glomerulus. (**C**) An adjacent H&E slice image of the brain region. (**D**) An adjacent H&E slice image of the kidney region. (**E–G**) Histogram distribution of the two regions in nucleic acid mass, protein mass, and quantum yield, respectively.

## References

1. J. D. Bancroft, M. Gamble, THEORY AND PRACTICE OF HISTOLOGICAL TECHNIQUES (Elsevier, ed. 8, 2018).

2. F. J. Fleming, A. D. K. Hill, E. W. Mc Dermott, A. O’Doherty, N. J. O’Higgins, C. M. Quinn, Intraoperative margin assessment and reexcision rate in breast conserving surgery. European Journal of Surgical Oncology (EJSO) 30, 233–237 (2004).

3. B. W. Maloney, D. M. McClatchy, B. W. Pogue, K. D. Paulsen, W. A. Wells, R. J. Barth, Review of methods for intraoperative margin detection for breast conserving surgery. J. Biomed. Opt. 23, 1 (2018).

4. W. H. Yong, Ed., Biobanking: Methods and Protocols (Springer New York, New York, NY, 2019; http://link.springer.com/10.1007/978-1-4939-8935-5)vol. 1897 of Methods in Molecular Biology.

5. F. Al-Mulla, Ed., Formalin-Fixed Paraffin-Embedded Tissues: Methods and Protocols (Humana Press, Totowa, NJ, 2011; https://link.springer.com/10.1007/978-1-61779-055-3)vol. 724 of Methods in Molecular Biology.

6. V. Rastogi, Artefacts: A Diagnostic Dilemma – A Review. JCDR, doi: 10.7860/JCDR/2013/6170.3541 (2013).

7. J. B. Taxy, Frozen Section and the Surgical Pathologist: A Point of View. Archives of Pathology & Laboratory Medicine 133, 1135–1138 (2009).

8. J. Kang, I. Song, H. Kim, H. Kim, S. Lee, Y. Choi, H. J. Chang, D. K. Sohn, H. Yoo, Rapid tissue histology using multichannel confocal fluorescence microscopy with focus tracking. Quant. Imaging Med. Surg 8, 884–893 (2018).

9. M. Rajadhyaksha, G. Menaker, T. Flotte, P. J. Dwyer, S. González, Confocal Examination of Nonmelanoma Cancers in Thick Skin Excisions to Potentially Guide Mohs Micrographic Surgery Without Frozen Histopathology. Journal of Investigative Dermatology 117, 1137–1143 (2001).

10. S. Abeytunge, B. Larson, G. Peterson, M. Morrow, M. Rajadhyaksha, M. P. Murray, Evaluation of breast tissue with confocal strip-mosaicking microscopy: a test approach emulating pathology-like examination. J. Biomed. Opt 22, 034002 (2017).

11. J. L. Dobbs, H. Ding, A. P. Benveniste, H. M. Kuerer, S. Krishnamurthy, W. Yang, R. Richards-Kortum, Feasibility of confocal fluorescence microscopy for real-time evaluation of neoplasia in fresh human breast tissue. J. Biomed. Opt 18, 106016 (2013).

12. E. Olson, M. J. Levene, R. Torres, Multiphoton microscopy with clearing for three dimensional histology of kidney biopsies. Biomed. Opt. Express 7, 3089 (2016).

13. Y. K. Tao, D. Shen, Y. Sheikine, O. O. Ahsen, H. H. Wang, D. B. Schmolze, N. B. Johnson, J. S. Brooker, A. E. Cable, J. L. Connolly, J. G. Fujimoto, Assessment of breast pathologies using nonlinear microscopy. Proc. Natl. Acad. Sci. U.S.A. 111, 15304–15309 (2014).

14. L. C. Cahill, M. G. Giacomelli, T. Yoshitake, H. Vardeh, B. E. Faulkner-Jones, J. L. Connolly, C.-K. Sun, J. G. Fujimoto, Rapid virtual hematoxylin and eosin histology of breast tissue specimens using a compact fluorescence nonlinear microscope. Laboratory Investigation 98, 150– 160 (2018).

15. M. Jain, B. D. Robinson, B. Wu, F. Khani, S. Mukherjee, Exploring Multiphoton Microscopy as a Novel Tool to Differentiate Chromophobe Renal Cell Carcinoma From Oncocytoma in Fixed Tissue Sections. Archives of Pathology & Laboratory Medicine 142, 383–390 (2018).

16. B. Sarri, F. Poizat, S. Heuke, J. Wojak, F. Franchi, F. Caillol, M. Giovannini, H. Rigneault, Stimulated Raman histology: one to one comparison with standard hematoxylin and eosin staining. Biomed. Opt. Express 10, 5378 (2019).

17. T. C. Hollon, B. Pandian, A. R. Adapa, E. Urias, A. V. Save, S. S. S. Khalsa, D. G. Eichberg, R. S. D’Amico, Z. U. Farooq, S. Lewis, P. D. Petridis, T. Marie, A. H. Shah, H. J. L. Garton, C. O. Maher, J. A. Heth, E. L. McKean, S. E. Sullivan, S. L. Hervey-Jumper, P. G. Patil, B. G. Thompson, O. Sagher, G. M. McKhann, R. J. Komotar, M. E. Ivan, M. Snuderl, M. L. Otten, T. D. Johnson, M. B. Sisti, J. N. Bruce, K. M. Muraszko, J. Trautman, C. W. Freudiger, P. Canoll, H. Lee, S. Camelo-Piragua, D. A. Orringer, Near real-time intraoperative brain tumor diagnosis using stimulated Raman histology and deep neural networks. Nat Med 26, 52–58 (2020).

18. D. A. Orringer, B. Pandian, Y. S. Niknafs, T. C. Hollon, J. Boyle, S. Lewis, M. Garrard, S. L. Hervey-Jumper, H. J. L. Garton, C. O. Maher, J. A. Heth, O. Sagher, D. A. Wilkinson, M. Snuderl, S. Venneti, S. H. Ramkissoon, K. A. McFadden, A. Fisher-Hubbard, A. P. Lieberman, T. D. Johnson, X. S. Xie, J. K. Trautman, C. W. Freudiger, S. Camelo-Piragua, Rapid intraoperative histology of unprocessed surgical specimens via fibrelaser-based stimulated Raman scattering microscopy. Nat Biomed Eng 1, 0027 (2017).

19. B. G. Saar, C. W. Freudiger, J. Reichman, C. M. Stanley, G. R. Holtom, X. S. Xie, Video-Rate Molecular Imaging in Vivo with Stimulated Raman Scattering. Science 330, 1368–1370 (2010).

20. A. Alfonso-García, R. Mittal, E. S. Lee, E. O. Potma, Biological imaging with coherent Raman scattering microscopy: a tutorial. J. Biomed. Opt 19, 071407 (2014).

21. P. Beard, Biomedical photoacoustic imaging. Interface Focus. 1, 602–631 (2011).

22. L. V. Wang, S. Hu, Photoacoustic Tomography: In Vivo Imaging from Organelles to Organs. Science 335, 1458–1462 (2012).

23. D.-K. Yao, Optimal ultraviolet wavelength for in vivo photoacoustic imaging of cell nuclei. J. Biomed. Opt 17, 056004 (2012).

24. J. Shi, T. T. W. Wong, Y. He, L. Li, R. Zhang, C. S. Yung, J. Hwang, K. Maslov, L. V. Wang, High-resolution, high-contrast mid-infrared imaging of fresh biological samples with ultraviolet-localized photoacoustic microscopy. Nat. Photonics 13, 609–615 (2019).

25. T. T. W. Wong, R. Zhang, P. Hai, C. Zhang, M. A. Pleitez, R. L. Aft, D. V. Novack, L. V. Wang, Fast label-free multilayered histology-like imaging of human breast cancer by photoacoustic microscopy. Sci. Adv. 3, e1602168 (2017).

26. N. J. M. Haven, K. L. Bell, P. Kedarisetti, J. D. Lewis, R. J. Zemp, Ultraviolet photoacoustic remote sensing microscopy. Opt. Lett. 44, 3586 (2019).

27. M. T. Martell, N. J. M. Haven, B. D. Cikaluk, B. S. Restall, E. A. McAlister, R. Mittal, B. A. Adam, N. Giannakopoulos, L. Peiris, S. Silverman, J. Deschenes, X. Li, R. J. Zemp, Deep learning-enabled realistic virtual histology with ultraviolet photoacoustic remote sensing microscopy. Nat Commun 14, 5967 (2023).

28. B. D. Cikaluk, B. S. Restall, N. J. M. Haven, M. T. Martell, E. A. McAlister, R. J. Zemp, Rapid ultraviolet photoacoustic remote sensing microscopy using voice-coil stage scanning. Opt. Express 31, 10136 (2023).

29. F. Fereidouni, Z. T. Harmany, M. Tian, A. Todd, J. A. Kintner, J. D. McPherson, A. D. Borowsky, J. Bishop, M. Lechpammer, S. G. Demos, R. Levenson, Microscopy with ultraviolet surface excitation for rapid slide-free histology. Nat Biomed Eng 1, 957–966 (2017).

30. T. C. Schlichenmeyer, M. Wang, K. N. Elfer, J. Q. Brown, Video-rate structured illumination microscopy for high-throughput imaging of large tissue areas. Biomed. Opt. Express 5, 366 (2014).

31. M. Wang, H. Z. Kimbrell, A. B. Sholl, D. B. Tulman, K. N. Elfer, T. C. Schlichenmeyer, B. R. Lee, M. Lacey, J. Q. Brown, High-Resolution Rapid Diagnostic Imaging of Whole Prostate Biopsies Using Video-Rate Fluorescence Structured Illumination Microscopy. Cancer Research 75, 4032–4041 (2015).

32. N. Kaza, A. Ojaghi, F. E. Robles, Virtual Staining, Segmentation, and Classification of Blood Smears for Label-Free Hematology Analysis. BME Front 2022, 9853606 (2022).

33. S. Soltani, A. Ojaghi, F. E. Robles, Deep UV dispersion and absorption spectroscopy of biomolecules. Biomed. Opt. Express 10, 487 (2019).

34. S. Soltani, A. Ojaghi, H. Qiao, N. Kaza, X. Li, Q. Dai, A. O. Osunkoya, F. E. Robles, Prostate cancer histopathology using label-free multispectral deep-UV microscopy quantifies phenotypes of tumor aggressiveness and enables multiple diagnostic virtual stains. Sci Rep 12, 9329 (2022).

35. S. Soltani, B. Cheng, A. O. Osunkoya, F. E. Robles, Deep UV Microscopy Identifies Prostatic Basal Cells: An Important Biomarker for Prostate Cancer Diagnostics. BME Front 2022, 9847962 (2022).

36. A. Ojaghi, E. Kendall Williams, N. Kaza, V. Gorti, H. Choi, J. Torey, T. Wiley, B. Turner, S. Jackson, S. Park, W. A. Lam, F. E. Robles, Label-free deep-UV microscopy detection and grading of neutropenia using a passive microfluidic device. Opt. Lett. 47, 6005 (2022).

37. V. Gorti, A. R. Subramanian, A. Ojaghi, J. Nsonwu-Farley, R. Tran, E. K. Williams, O. Torres, A. Aljudi, W. Aumann, F. E. Robles, Rapid, Point-of-Care Bone Marrow Aspirate Adequacy Assessment Via Deep Ultraviolet Microscopy. Laboratory Investigation 105, 104102 (2025).

38. V. Gorti, K. McCubbins, D. Houston, A. D. Silva Trenkle, A. Holberton, C. E. Serafini, L. Wood, G. Kwong, F. E. Robles, Quantifying UV-induced photodamage for longitudinal live-cell imaging applications of deep-UV microscopy. Biomed. Opt. Express 16, 208 (2025).

39. V. Gorti, N. Kaza, E. K. Williams, W. A. Lam, F. E. Robles, Compact and low-cost deep-ultraviolet microscope system for label-free molecular imaging and point-of-care hematological analysis. Biomed. Opt. Express 14, 1245 (2023).

40. A. Ojaghi, M. E. Fay, W. A. Lam, F. E. Robles, Ultraviolet Hyperspectral Interferometric Microscopy. Sci Rep 8, 9913 (2018).

41. B. J. Zeskind, C. D. Jordan, W. Timp, L. Trapani, G. Waller, V. Horodincu, D. J. Ehrlich, P. Matsudaira, Nucleic acid and protein mass mapping by live-cell deep-ultraviolet microscopy. Nat Methods 4, 567–569 (2007).

42. S. Ye, J. Zou, C. Huang, F. Xiang, Z. Wen, N. Wang, J. Yu, Y. He, P. Liu, X. Mei, H. Li, L. Niu, P. Gong, W. Zheng, Rapid and label-free histological imaging of unprocessed surgical tissues via dark-field reflectance ultraviolet microscopy. iScience 26, 105849 (2023).

43. Y. Kumamoto, A. Taguchi, S. Kawata, Deep-Ultraviolet Biomolecular Imaging and Analysis. Advanced Optical Materials 7, 1801099 (2019).

44. Chunqiang Li, C. Pitsillides, J. M. Runnels, D. Côté, C. P. Lin, Multiphoton Microscopy of Live Tissues With Ultraviolet Autofluorescence. IEEE J. Select. Topics Quantum Electron. 16, 516–523 (2010).

45. J. R. Lakowicz, B. R. Masters, Principles of Fluorescence Spectroscopy, Third Edition. J. Biomed. Opt. 13, 029901 (2008).

46. H. Liu, H. Zhang, B. Jin, Fluorescence of tryptophan in aqueous solution. Spectrochimica Acta Part A: Molecular and Biomolecular Spectroscopy 106, 54–59 (2013).

47. W. F. Cheong, S. A. Prahl, A. J. Welch, A review of the optical properties of biological tissues. IEEE J. Quantum Electron. 26, 2166–2185 (1990).

48. J. Ritz, A. Roggan, C. Isbert, G. Müller, H. J. Buhr, C. Germer, Optical properties of native and coagulated porcine liver tissue between 400 and 2400 nm. Lasers Surg Med 29, 205–212 (2001).

49. K. Tsia, Ed., Understanding Biophotonics: Fundamentals, Advances, and Applications (Jenny Stanford Publishing, ed. 0, 2016; https://www.taylorfrancis.com/books/9789814411783).

50. I. S. Martins, H. F. Silva, E. N. Lazareva, N. V. Chernomyrdin, K. I. Zaytsev, L. M. Oliveira, V. V. Tuchin, Measurement of tissue optical properties in a wide spectral range: a review [Invited]. Biomed. Opt. Express 14, 249 (2023).

51. M. S. Nogueira, S. Maryam, M. Amissah, H. Lu, N. Lynch, S. Killeen, M. O’Riordain, S. Andersson-Engels, Evaluation of wavelength ranges and tissue depth probed by diffuse reflectance spectroscopy for colorectal cancer detection. Sci Rep 11, 798 (2021).

52. M. N. S. Yusoff, M. S. Jaafar, EFFECT OF MULTI-DESIGN SKIN MODEL AND CHARACTERISTIC ON MONTE CARLO SIMULATION OF LIGHT-SKIN DIFFUSE REFLECTANCE SPECTRA. Jurnal Teknologi 78 (2016).

53. E. L. Wisotzky, F. C. Uecker, S. Dommerich, A. Hilsmann, P. Eisert, P. Arens, Determination of optical properties of human tissues obtained from parotidectomy in the spectral range of 250 to 800 nm. J. Biomed. Opt. 24, 1 (2019).

54. T. M. Gonçalves, I. S. Martins, H. F. Silva, V. V. Tuchin, L. M. Oliveira, Spectral Optical Properties of Rabbit Brain Cortex between 200 and 1000 nm. Photochem 1, 190–208 (2021).

55. T. Yoshitake, M. G. Giacomelli, L. M. Quintana, H. Vardeh, L. C. Cahill, B. E. Faulkner-Jones, J. L. Connolly, D. Do, J. G. Fujimoto, Rapid histopathological imaging of skin and breast cancer surgical specimens using immersion microscopy with ultraviolet surface excitation. Sci Rep 8, 4476 (2018).

56. R. Sidman, J. Angevine, E. T. Pierce, High Resolution Mouse Brain Atlas - Methods.

57. T. Tekko, T. Lakspere, A. Allikalt, J. End, K. R. Kõlvart, T. Jagomäe, A. Terasmaa, M.-A. Philips, T. Visnapuu, F. Väärtnõu, S. F. Gilbert, A. Rinken, E. Vasar, K. Lilleväli, Wfs1 is expressed in dopaminoceptive regions of the amniote brain and modulates levels of D1-like receptors. PLoS ONE 12, e0172825 (2017).

58. S. Suzuki, A. Mori, Regional distribution of tyrosine, tryptophan, and their metabolites in the brain of epileptic El mice. Neurochem Res 17, 693–698 (1992).

59. A. C. Ruifrok, D. A. Johnston, Quantification of histochemical staining by color deconvolution. Anal Quant Cytol Histol 23, 291–299 (2001).

60. T. V. Waehneldt, E. M. Shooter, A comparison of the protein composition of the brains of four rodents. Brain Research 57, 361–371 (1973).

61. M. K. Fernandez, M. Sinha, M. Renz, Differential Intracellular Protein Distribution in Cancer and Normal Cells—Beta-Catenin and CapG in Gynecologic Malignancies. Cancers 14, 4788 (2022).

62. J. Li, J. Garfinkel, X. Zhang, D. Wu, Y. Zhang, K. De Haan, H. Wang, T. Liu, B. Bai, Y. Rivenson, G. Rubinstein, P. O. Scumpia, A. Ozcan, Biopsy-free in vivo virtual histology of skin using deep learning. Light Sci Appl 10, 233 (2021).

63. Y. Zhang, L. Kang, I. H. M. Wong, W. Dai, X. Li, R. C. K. Chan, M. K. Y. Hsin, T. T. W. Wong, High-Throughput, Label-Free and Slide-Free Histological Imaging by Computational Microscopy and Unsupervised Learning. Advanced Science 9, 2102358 (2022).

64. M. Boktor, J. E. D. Tweel, B. R. Ecclestone, J. A. Ye, P. Fieguth, P. Haji Reza, Multi-channel feature extraction for virtual histological staining of photon absorption remote sensing images. Sci Rep 14, 2009 (2024).

65. B. R. Ecclestone, J. A. T. Simmons, J. E. D. Tweel, C. Kaur, A. Hajiahmadi, P. Haji, Photon Absorption Remote Sensing (PARS): A Comprehensive Approach to Label-free Absorption Microscopy Across Biological Scales. [Preprint] (2024). 10.48550/arXiv.2403.04229.

66. J.-Y. Zhu, T. Park, P. Isola, A. A. Efros, “Unpaired Image-to-Image Translation Using Cycle-Consistent Adversarial Networks” in 2017 IEEE International Conference on Computer Vision (ICCV) (IEEE, Venice, 2017; http://ieeexplore.ieee.org/document/8237506/), pp. 2242–2251.

67. T. M. Abraham, P. C. Costa, C. E. Serafini, Z. Guang, Z. Zhang, S. Neill, J. J. Olson, R. Levenson, F. E. Robles, Label- and slide-free tissue histology using 3D epi-mode quantitative phase imaging and virtual hematoxylin and eosin staining. Optica 10, 1605 (2023).

68. P. Ledwig, F. E. Robles, Epi-mode tomographic quantitative phase imaging in thick scattering samples. Biomed. Opt. Express 10, 3605 (2019).

69. P. Ledwig, F. E. Robles, Quantitative 3D refractive index tomography of opaque samples in epi-mode. Optica 8, 6 (2021).

70. P. Ledwig, M. Sghayyer, J. Kurtzberg, F. E. Robles, Dual-wavelength oblique back-illumination microscopy for the non-invasive imaging and quantification of blood in collection and storage bags. Biomed. Opt. Express 9, 2743 (2018).

71. P. C. Costa, Z. Guang, P. Ledwig, Z. Zhang, S. Neill, J. J. Olson, F. E. Robles, Towards in-vivo label-free detection of brain tumor margins with epi-illumination tomographic quantitative phase imaging. Biomed. Opt. Express 12, 1621 (2021).

72. P. Casteleiro Costa, B. Wang, C. Filan, A. Bowles-Welch, C. Yeago, K. Roy, F. E. Robles, Functional imaging with dynamic quantitative oblique back-illumination microscopy. J. Biomed. Opt. 27 (2022).

73. E. Angioni, G. Lercker, N. G. Frega, G. Carta, M. P. Melis, E. Murru, S. Spada, S. Banni, UV spectral properties of lipids as a tool for their identification. Eur. J. Lipid Sci. Technol. 104, 59–64 (2002).

74. Z. Li, P. Casteleiro Costa, C. Serafini, S. Bharadwaj, Z. Guang, F. E. Robles, Experimental assessment of the optical transfer function for quantitative oblique back illumination microscopy (qOBM). Opt. Express 33, 5088 (2025).

75. T. Zheng, W. Adi, P. J. Campagnola, F. Yesilkoy, Fix or Freeze? Spectral Differences Arising from Tissue Preparation in Chemical Imaging. [Preprint] (2025). 10.1101/2025.11.19.689284.

76. T. N. Ford, K. K. Chu, J. Mertz, Phase-gradient microscopy in thick tissue with oblique back-illumination. Nat Methods 9, 1195–1197 (2012).

77. Z. Li, P. Casteleiro Costa, Z. Guang, C. E. Serafini, F. E. Robles, GAN-based quantitative oblique back-illumination microscopy enables computationally efficient epi-mode refractive index tomography. Biomed. Opt. Express 15, 4764 (2024).

78. B. J. Zeskind, C. D. Jordan, W. Timp, L. Trapani, G. Waller, V. Horodincu, D. J. Ehrlich, P. Matsudaira, Nucleic acid and protein mass mapping supp. Nat Methods 4, 567–569 (2007).

79. T. Li, C. Zhou, M. Jiang, UV absorption spectra of polystyrene. Polymer Bulletin 25, 211–216 (1991).

80. M. Meinhardt, R. Krebs, A. Anders, U. Heinrich, H. Tronnier, Wavelength-dependent penetration depths of ultraviolet radiation in human skin. J. Biomed. Opt. 13, 044030 (2008).

